# Encephalitis patient derived monoclonal GABA_A_ receptor antibodies cause catatonia and epileptic seizures

**DOI:** 10.1101/2021.01.28.428602

**Authors:** Jakob Kreye, Sukhvir K. Wright, Adriana van Casteren, Marie-Luise Machule, S. Momsen Reincke, Marc Nikolaus, Laura Stöffler, Scott van Hoof, Elisa Sanchez-Sendin, Hans-Christian Kornau, Angela M. Kaindl, Max A. Wilson, Stuart Greenhill, Gavin Woodhall, Paul Turko, Imre Vida, Craig C. Garner, Jonathan Wickel, Christian Geis, Yuko Fukata, Masaki Fukata, Harald Prüss

## Abstract

Autoantibodies targeting the GABA_A_ receptor (GABA_A_R) hallmark an autoimmune encephalitis presenting with frequent seizures and psychomotor abnormalities. Their pathogenic role is still not well-defined, given the common overlap with further autoantibodies and the lack of patient derived monoclonal antibodies (mAbs). We cloned and recombinantly produced five affinity-maturated GABA_A_R IgG1 mAbs from cerebrospinal fluid cells, which bound to various epitopes involving α1 and γ2 receptor subunits, with variable binding strength and partial competition. mAbs selectively reduced GABAergic currents in neuronal cultures without causing receptor internalization. Cerebroventricular infusion of GABA_A_R mAbs and Fab fragments into rodents induced a severe phenotype with catatonia, seizures and increased mortality, reminiscent of encephalitis patients’ symptoms. Our results prove direct functional effects of autoantibodies on GABA_A_Rs and provide an animal model for GABA_A_R encephalitis. They further provide the scientific rationale for clinical treatments using antibody depletion and pave the way for future antibody-selective immunotherapies.

## Introduction

GABA_A_ receptors (GABA_A_Rs) are key molecules for physiological brain function, transmitting rapid phasic inhibitory synaptic signaling and mediating tonic inhibition at extrasynaptic and perisynaptic locations^1^. The pentameric ligand gated chloride channels can be composed of different subunits (α1–6, β1–3, γ1–3, δ, ε, π, θ, and σ1–3), most abundantly in the α1β2γ2 configuration^2^. Receptor dysfunction can lead to severe neurological symptoms, such as epileptic encephalopathies based on GABA_A_R subunit mutations3-6. Recently, cerebrospinal fluid (CSF) and serum autoantibodies targeting the α1, β3 and γ2 subunits of GABA_A_Rs were identified in a new form of autoimmune encephalitis presenting with seizures, refractory status epilepticus, cognitive alterations, psychomotor disorders and MRI abnormalities^7–10^. Patients’ sera or CSF containing polyclonal GABA_A_R antibodies caused downregulation of surface GABA_A_R and electrophysiological changes in cultured neurons^7–9.^

Patients with GABA_A_R encephalitis frequently harbor further established pathogenic autoantibodies such as those targeting LGI1, CASPR2, NMDAR^7–9^, hence it is unclear whether the observed effects exclusively relate to GABA_A_R antibodies. Interestingly, in a subset of patients antibodies against intracellular GAD65 were observed^8^ and recently strongly expanded CD8+ T cell clones have been described^11^, both pointing towards an accompanying T cell driven immune response.

In this current study, we aimed to characterize the intrathecal human monoclonal antibody (mAb) repertoire from antibody-secreting cells (ASCs) and B cells from cerebrospinal fluid in acute GABA_A_R encephalitis. Using recombinant production of CSF derived mAbs12, 13, we generated a set of GABA_A_R mAbs for the characterization of antibody sequence features, epitope mapping and pathogenic functional effects *in vitro* and *in vivo*, independent of confounding factors.

## Results

### Monoclonal CSF antibodies from encephalitis patient target GABAAR and non- GABA_A_R antigens

To investigate the functional role of GABA_A_R antibodies in encephalitis pathogenesis, we first explored the monoclonal immunoglobulin repertoire in the CSF of a pediatric GABA_A_R encephalitis patient presenting with catatonia^14^. The antibody response was captured from single cells of three populations: CD138^+^ ASCs, CD20^+^CD27^+^ memory B cells (MBCs) and CD20^+^CD27^−^ non-memory B cells (NMBCs), separated via fluorescence-activated cell sorting (Supplementary Fig. 1). Using single cell cloning^12, 15^, we generated 67 recombinant human mAbs, which were screened for GABA_A_R reactivity on cell-based assays (CBAs) and on unfixed murine whole brain sections as an unbiased test for central nervous system (CNS) auto-reactivity.

We identified five different human GABA_A_R mAbs. Four reacted positive in two CBAs using human embryonic kidney (HEK) cells expressing either α1β3 (comparable to clinical routine assays) or α1β3γ2 GABA_A_Rs (Supplementary Fig. 2). In contrast, mAb #113-175 bound to GABA_A_Rs on the α1β3γ2 CBA only (Supplementary Fig. 2). All five mAbs revealed strong tissue reactivity on murine brain sections (Fig. 1a-c, Supplementary Fig. 3), most prominent against hippocampal neuropil (Fig. 1a, b), the granule cell and molecular layer in the cerebellum (Fig. 1c), putamen and olfactory bulb. Reactivity was confirmed on cultured rat neurons and showed clustered distribution of GABA_A_Rs along MAP2+ dendrites (Fig. 1d), overlapping with a commercial rabbit GABA_A_R-α1 antibody and co-localizing with presynaptic vesicular GABA transporter (VGAT, Fig. 1e). In addition to typical neuronal GABA_A_R binding pattern, #113-201 revealed intense reactivity against choroid plexus and around blood vessels (Supplementary Fig. 3g-i).

**Fig. 1.**
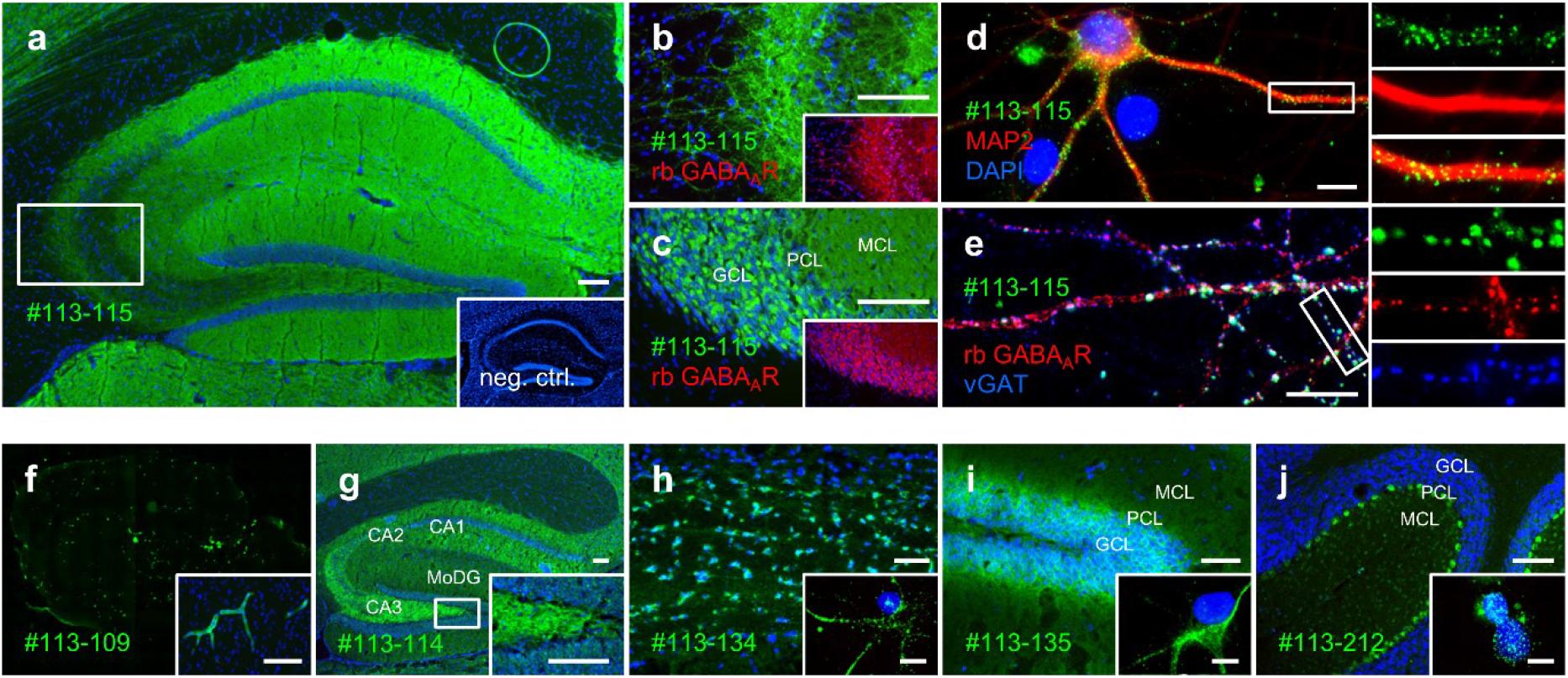
Binding of human monoclonal antibodies to GABA_A_R and other CNS targets. (a-c) Immunofluorescence stainings of human GABA_A_R mAb #113-115 (green) on unfixed murine brain tissue; with binding to (a-b) hippocampal neuropil and (c) in the granule cell (GCL) and molecular (MCL) but not Purkinje cell (PCL) layer of the cerebellum, in co-localization with a commercial anti-GABA_A_R antibody (red, nuclei in blue). (d-e) Immunofluorescence staining on cultured cortical rat neurons, with GABA_A_R mAb #113-115 (green) in (d) clustered binding pattern along MAP2^+^ (red) dendrites and in (e) co-localization with commercial antibodies against GABA_A_R (red) and the pre-synaptic marker vesicular GABA transporter (vGAT, blue). (f-j) Immunofluorescence staining of human GABA_A_R - negative mAbs (green, nuclei in blue) from encephalitis patient’s CSF repertoire with specific brain tissue reactivity in variable distribution patterns, including binding to (f) blood vessels, (g) hippocampal neuropil, (h) cell soma in corpus callosum and (i-j) cells of different cerebellar layers. Inserts in (h-j) show mAb staining on cultured neurons. Scale bars indicate 100 μm in (a-c), 20 μm in (d-e), 100 μm in (f-j) and 20 μm in inserts of (f-j).

Seventeen of the expressed GABA_A_R-negative mAbs (27.9%) showed intense tissue binding on murine brain sections in distinct patterns, mostly on neuronal surfaces of the hippocampus, cerebellum and in basal ganglia, but others also against blood vessels, choroid plexus, ependyma and white matter tracts (Fig. 1f-j, Supplementary Fig. 4). A subset also reacted with cultured neurons (Fig. 1h-j inserts). Screening for already established neuronal antigens with commercial assays identified Homer-3 as the target of #113-212 (Fig. 1j), while the other non-GABA_A_R antigens remained unknown.

### Polyclonal response to GABA_A_Rs derived mainly from IgG1 antibody secreting cells

We next investigated sequence features of the GABA_A_R encephalitis immunoglobulin repertoire. mAbs derived from different cell populations revealed characteristic Ig isotype distributions (Fig. 2a, Supplementary Table 1). All GABA_A_R mAbs were of the IgG1 subtype (Fig. 2b) and derived predominantly from ASCs, also from one MBC, but not from NMBCs (Fig. 2c). In contrast, CNS tissue-reactive, but GABA_A_R-negative mAbs were similarly distributed within different cell populations (Supplementary Table 1). Among all mAb sequences no clonal relationships were identified, indicating a polyclonal response to GABA_A_Rs.

**Fig. 2.**
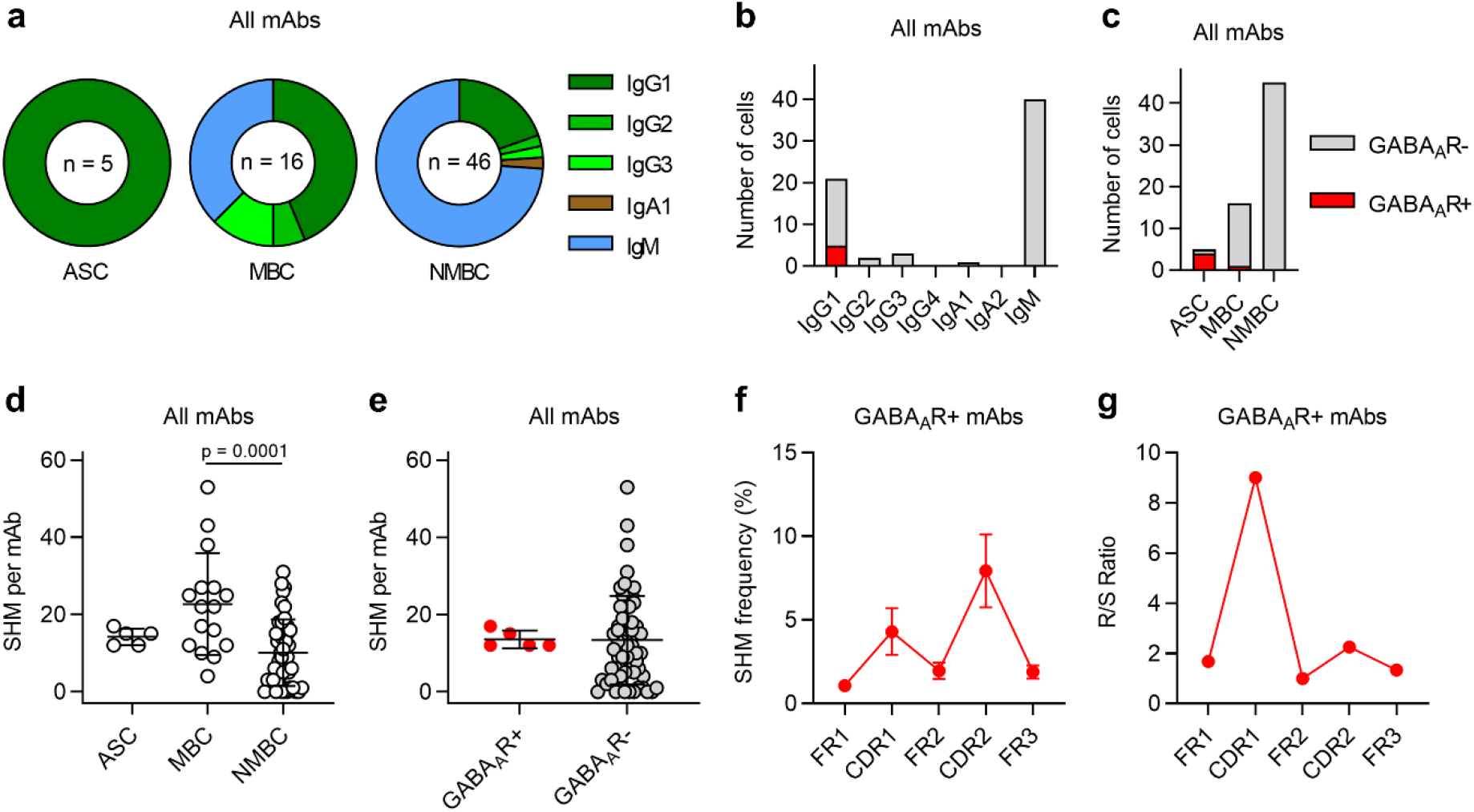
Ig sequence data analysis from GABA_A_R encephalitis CSF repertoire. (a) Ig subclass distributions per mAb source cell type from GABA_A_R encephalitis CSF repertoire. (b-c) Absolute frequencies of GABA_A_R-reactive (GABA_A_R+) and GABA_A_R-negative (GABA_A_R-) mAbs per (b) Ig subclass and (c) mAb source cell type. (d) Comparison of SHM counts in the variable domain V genes between mAbs of different source cell types (ASC n = 5, MBC n = 16, NMBC = 46). Data was analyzed using ordinary one-way ANOVA followed by posthoc Tukey’s multiple comparisons tests (p values not shown when not significant). Bars indicate mean ± SD. (e) Comparison of SHM counts in the variable domain V genes between GABA_A_R+ (n = 5) and GABA_A_R-(n = 62) mAbs. Bars indicate mean ± SD. (f) Relative frequencies of SHM per nucleotide within complementarity-determining regions (CDR) and frame regions (FR) of GABA_A_R+ mAb genes, shown as mean ± SEM, n = 5. (g) Ratios of replacement to silence (R/S) mutations within CDRs and FRs for all SHM of all GABA_A_R+ mAb genes combined.

Variable domains from MBC derived mAbs had more somatic hypermutations (SHMs) than NMBC derived mAbs on heavy chains, light chains and per mAb in total (22.6 ± 12.8 vs. 10.1 ± 8.6; p=0.0001, ANOVA, posthoc Tukey’s multiple comparisons test; Fig. 2d, Supplementary Table 1), reflecting affinity maturation. SHM counts of ASC mAbs (14.2 ± 1.9) were between those of MBC and NMBC mAbs (Fig. 2d). GABA_A_R mAbs contained a similar number of SHMs as GABA_A_R-negative mAbs (Fig. 2e). Within GABA_A_R mAbs SHM were higher in frequency and replacement to silence ratios in complementarity-determining regions (CDR) than in frame regions (FR) (Fig. 2f-g), characteristic for antigen driven maturation.

### Patient derived mAbs bound α1 and γ2 subunits of GABA_A_R with different strength and partial competition

To select the most relevant antibodies for functional studies we aimed to characterize the subunit specificity and the binding strength of the isolated GABA_A_Rs mAbs. To this end, we first performed a series of CBAs expressing individual GABA_A_R subunits or combinations thereof. The patient’s polyclonal antibodies from CSF and plasmapheresis eluate (PPE) showed predominantly α1 reactivity and additionally bound very weakly to COS7 cells expressing α2β3, α5β3 and β3γ2 subunit combinations (PPE only) (Fig. 3a, Supplementary Fig. 5a). mAb stainings confirmed the α1 subunit as the main target from GABA_A_R mAbs (Fig. 3a) and excluded binding to α1-5, β1-3 or γ2 subunits for all further mAbs with anti-neuronal reactivity (Fig. 3a, exemplarily shown for #113-109). In detail, #113-101, #113-115 and #113-198 selectively target α1, as these antibodies detected COS7 cells expressing α1β3 and α1β3γ2, but not β3 nor γ2 alone (Fig. 3a). Specificity was confirmed by the absence of reactivity to α2β3γ2, α3β3γ2, α4β3γ2 and α5β3γ2 (not shown). #113-201 also targeted α1, as it bound to α1β3, but not β3 alone and also not to α2β3, α3β3, α4β3 and α5β3 (Fig. 3a, Supplementary Fig. 5b). Additionally, #113-201 detected γ2 alone (Fig. 3a), which has about 45% identity with the α1 subunit. Interestingly, #113-175 bound to α1β3γ2, but neither to α1β3, β3 alone, γ2 alone nor to β3γ2 (Fig. 3a), suggesting that #113-175 recognizes a three-dimensional epitope of GABA_A_R heteromers requiring the co-expression of α1 and γ2. Consistently, #113-175 reacted with α1β2γ2, but not with α2β3γ2, α3β3γ2, α4β3γ2 and α5β3γ2 (Supplementary Fig. 5c).

**Fig. 3.**
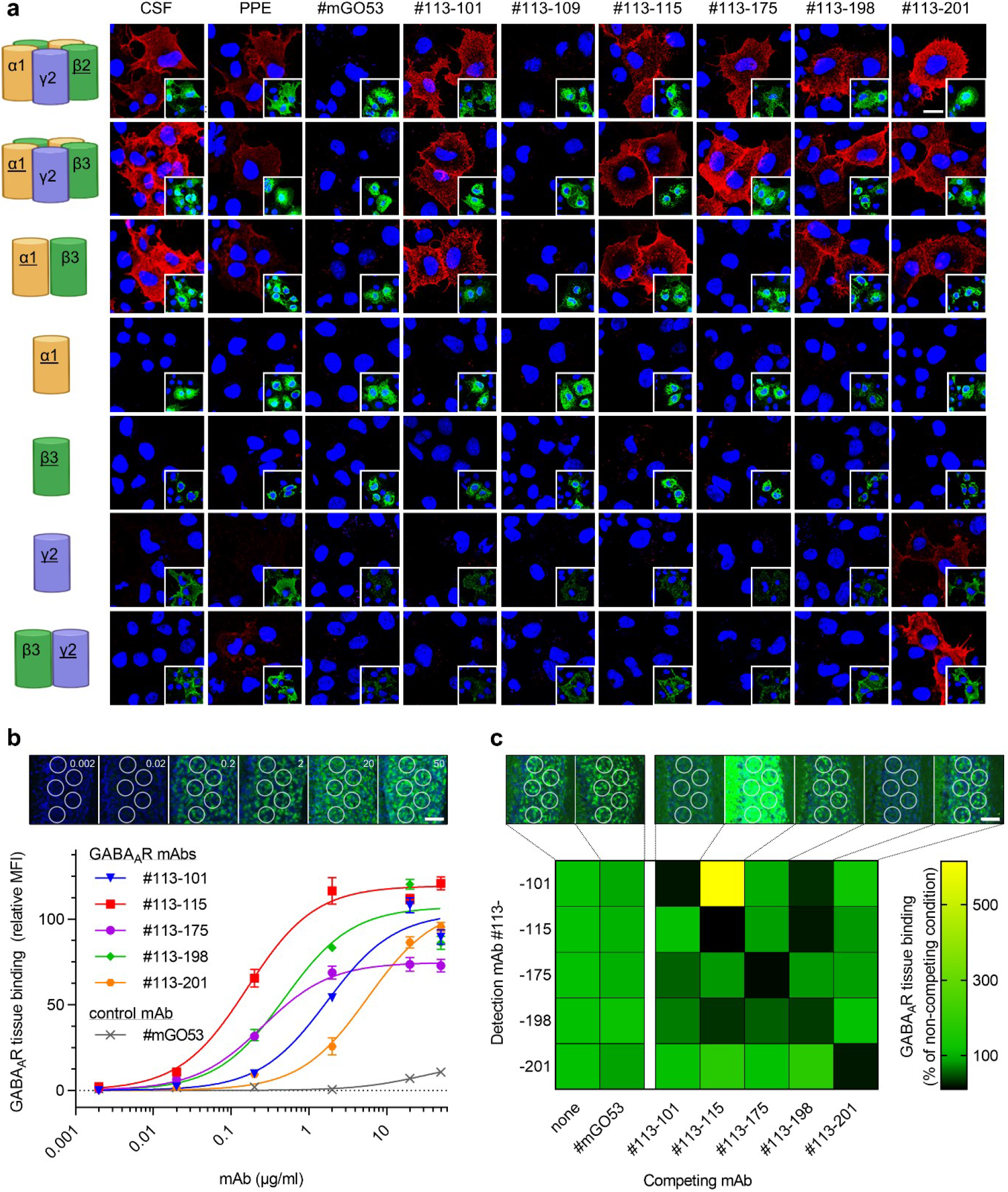
GABA_A_R subunit epitopes and mAb binding properties. (a) Immunofluorescence stainings as cell-based assays (CBA) using COS7 cells overexpressing individual or multiple GABA_A_R subunits (as illustrated in left column) to evaluate subunit-specific binding of patient’s polyclonal samples and derived recombinant mAbs (red, nuclei in blue). Negative controls #mGO53 and #113-109 showed no binding. Underlined subunits were stained with subunit-specific commercial antibodies (shown in green in image inserts). Note that β3 alone is expressed on the cell-surface, but α1 alone is not^38^. Representative scale bar indicates 20 μm. (b) For relative quantification of human GABA_A_R mAb binding to natively expressed receptors the mean fluorescence intensity (MFI) was measured from binding in the granule cell layer of the murine cerebellum. Representative images from #113-115 are shown in top row with IgG concentrations as indicated. Representative scale bar indicates 100 μm. MFIs from 15 regions of interest (ROI) from 3 independent experiments were used to fit non-linear regression models of specific binding. Error bars indicate SEM. (c) For analysis of competitive binding, fluorophore-coupled GABA_A_R mAbs (detection mAbs) were stained on murine brain tissue in combination with of an uncoupled mAb (competing mAb) in excess. Exemplary images from coupled #113-101 are shown in top row with a representative scale bar indicating 100 μm. Quantified mean MFIs as relative values to non-competing condition of the respective coupled mAb are shown as a heat map, each derived from 45 ROI from 3 independent experiments. Receptor binding competition is visualized in black and signal enhancement in yellow.

For the identification of the mAb with strongest binding to natively expressed GABA_A_Rs we then quantified binding of purified mAbs at serial dilutions to murine cerebellum (Fig. 3b). Binding curves were derived from mean fluorescent intensities (MFI) to quantify maximum intensities (MFI_max_) and concentrations at which 50% of MFI_max_ is reached (Half Max; Supplementary Table 2). #113-115 showed strongest reactivity (Fig. 3b, red) as indicated by lowest Half Max of 160 ng/ml, which was approximately 40-fold higher than from the weakest binder #113-201 (Fig. 3b, orange). Models from four out of five mAbs displayed similar MFI_max_. This indicates likewise similar available binding sites, which is for α1-selective mAbs represented by the presence of two α1 subunits and therefore two binding sites per heteropentameric GABA_A_R. In contrast, the target epitope of #113-175 involves the α1 and γ2 subunit, therefore can likely be present only once per GABA_A_R. Consistently, mAb #113-175 reached a considerably lower MFI_max_, but with a similar concentration dependency as #113-115 (Fig 3b, purple). In a complementary approach using flow cytometry, we found similar mAb binding to HEK cells overexpressing α1β3γ2 GABA_A_R subunits, thus excluding potential bias related to unspecific tissue binding (Supplementary Fig. 6a-d).

As GABA_A_R binding for all mAbs involves α1, we analyzed whether target epitopes are identical or in close proximity in a competition assay. GABA_A_R mAbs were fluorophore-coupled and used for quantitative detection of binding to murine cerebellum. The binding of each fluorophore-coupled GABA_A_R mAb was abrogated in the presence of its respective unlabeled mAb at excess, but not when co-applied with non-reactive control mAb #mGO53 (Fig. 3c). Quantification of all possible pairings of GABA_A_R mAbs revealed that certain mAbs can decrease the binding of other GABA_A_R mAbs (Fig. 3c, black tiles), suggesting competition for overlapping target epitopes. In contrast, #113-201 did not influence receptor binding of any other GABA_A_R mAb. Interestingly, the binding signal of #113-101 was markedly increased in the presence of unlabeled #113-115 (Fig. 3c, yellow tile) and similarly with #113-115 Fab fragments (Supplementary Fig. 7a). As #113-101 did not directly bind #113-115 (Supplementary Fig. 7b, c), this finding suggests conformational GABA_A_R changes induced by #113-115 binding, leading to increased #113-101 epitope accessibility.

### mAb #113-115 selectively reduced GABAergic currents without causing receptor internalization

Next, we explored the pathogenic relevance of single GABA_A_R mAbs on GABAergic functions *in vitro*. We selected #113-115 being the mAb with strongest binding to native GABA_A_Rs and #113-175 which depends on α1γ2-co-expression. In electrophysiological recordings from cultured autaptic neurons incubated for 24 hours with 1 μg/ml of #113-175 we observed no differences to untreated and control conditions. In contrast, #113-115 led to reduced inhibitory postsynaptic signaling as indicated in lower amplitudes of evoked IPSCs (Fig. 4a, c) in comparison to untreated condition and to control mAb treatment (#113-115: 0.31 ± 0.09 SEM; untreated: 1 ± 0.12, p < 0.0001, Kruskal-Wallis, Dunn’s posthoc; control: 0.91 ± 0.16, p = 0.0037). This effect was GABA-specific, as amplitudes of selective responses to GABA (Fig. 4b, d) were likewise reduced (#113-115: 0.55 ± 0.06; untreated: 1 ± 0.09, p = 0.0011; control: 1.02 ± 0.07, p = 0.0001), whereas responses to kainate and NMDA remained unaffected (Fig. 4e, f).

**Fig. 4.**
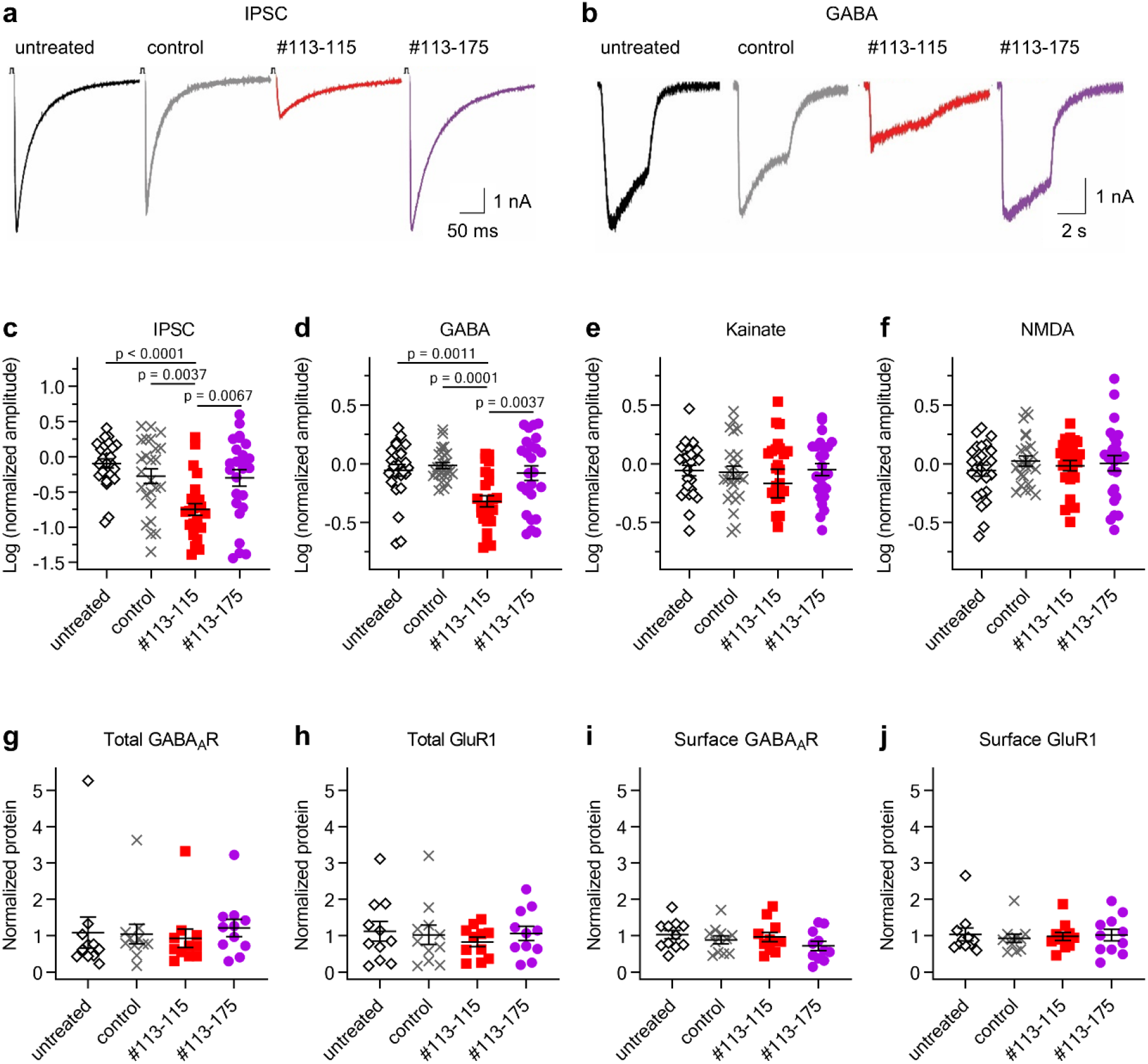
mAb #113-115 selectively reduced GABAergic signalling *in vitro*, independent of receptor internalization. (a-b) Representative traces from (a) evoked or (b) GABA-dependent currents of murine autaptic neurons after pre-incubation with indicated human mAb. (c-f) Amplitudes of (c) evoked or chemically induced responses to (d) GABA, (e) kainate or (f) NMDA of murine autaptic neurons after mAb pre-incubation as indicated (values normalized to mean of untreated condition, n = 25 per condition). Data was analyzed using Kruskal-Wallis, Dunn’s posthoc tests (p values not shown when not significant). (g-j) Quantifications of the indicated (g-h) total and and (i-j) biotinylated surface proteins from cultured neocortical rat neurons as analyzed by Western blotting and normalized to N-Cadherin expression and mean of untreated condition. Error bars indicate SEM; n = 4 separate cultures.

Previous studies using patients’ sera or CSF containing GABA_A_R antibodies suggested receptor internalization as a pathogenic mechanism in GABA_A_R encephalitis. To examine this mechanism at the level of mAbs, we quantified protein levels from neuronal cultures after mAb incubation (Supplementary Fig. 8a-c, Fig. 4g-j). The expression levels of GABA_A_R and control protein GluR1 were unaltered under mAb treatment and control conditions, both in the total protein (Fig. 4g-h) and in the surface protein fractions (Fig. 4i-j). Surface to total protein ratios of the intracellular marker mortalin were similar in all groups, indicating no mAb influence on cell membrane integrity and thus cell viability (Supplementary Fig. 8d). Control experiments assured the presence and stability of mAbs after the culture conditions (Supplementary Fig. 8e, f). These results were independently confirmed using reader-based immunocytochemistry to quantify surface GABA_A_R levels of neuronal cultures, which displayed no changes between all groups (Supplementary Fig. 8g-h).

### Cerebroventricular infusion of #113-115 IgG and Fab fragments induced encephalopathic symptoms and increased mortality

The effects observed *in vitro* indicate direct pathogenicity of antibody-antigen binding, independent of receptor internalization, a mechanism related to bivalent antibody mediated crosslinking^16^. Thus, we next assessed whether mAbs can similarly induce pathogenic effects *in vivo* and applied GABA_A_R mAb #113-115 in mice by continuous cerebroventricular infusion via osmotic pumps. To also explore the role of the antibody’s bivalence we generated and characterized a monovalent Fab fragment version of #113-115 lacking the Fc domain (Supplementary Fig. 9a, b).

Within a few days after implantation, six out of seven mice of the #113-115 IgG high-dose group and all five mice of the #113-115 Fab high-dose group developed encephalopathic symptoms compatible with impaired GABAergic inhibition, including myoclonus, twitching, gait ataxia and circling (Supplementary Video 1). In the #113-115 IgG low-dose group only two out of six mice developed similar symptoms which started later after pump implantation. No disease symptoms were observed in the control mice after mAb #mGO53 infusion. All symptomatic mice in the #113-115 IgG high-dose and #113-115 Fab group died or had to be sacrificed after reaching predefined humane endpoints, commonly status epilepticus (Fig. 5d). Extracted brains from #113-115-infused but not #mGO53-infused mice revealed intense deposition of human IgG or Fab in characteristic GABA_A_R distribution pattern (Fig. 5a-c), indicating *in vivo* antigen binding. However, the levels of GABA_A_R and control proteins from homogenized extracted brains showed no differences between the groups (Fig. 5e-h), consistent with the *in vitro* data (Fig. 4g-j, Supplementary Fig. 8g-h).

**Fig. 5.**
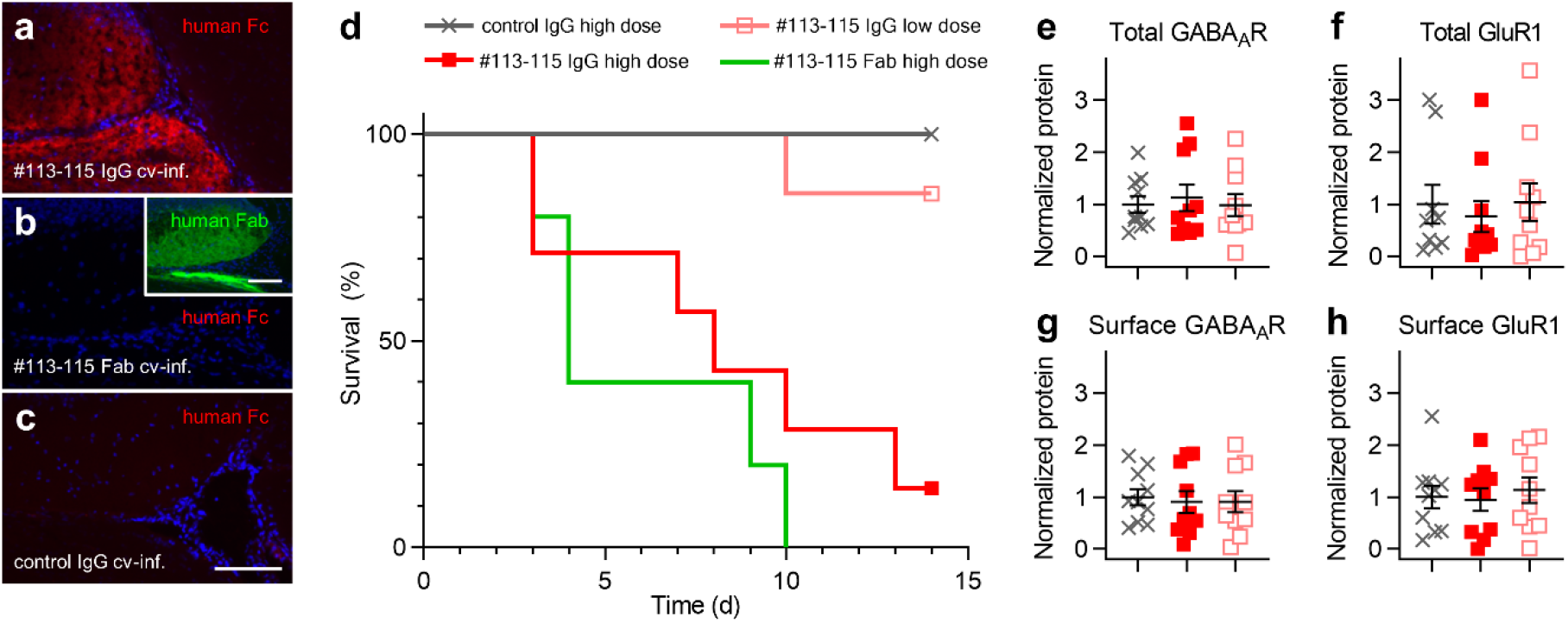
GABA_A_R IgG and Fab induced increased mortality. (a-c) Immunofluorescence stainings on brain sections from C57BL/6 mice after cerebroventricular infusion (cv-inf.) of the indicated GABA_A_R or control mAbs over 14 days. Detection with anti-human Fc-specific antibody (red) or anti-human Fab-specifc antibody (green, insert in b) revealed characteristic hippocampal mAb deposition of (a) #113-115 IgG and (b) #113-115 Fab; (c) not seen in control animals receiving control IgG (#mGO53). Representative scale bars indicate 100 μm. (d) Kaplan-Meier plot for survival of C57BL/6 mice after cerebroventricular infusion over 14 days of indicated mAbs as IgG in high or low dose (1.5/0.3 μg IgG per hour) or as Fab in high dose. Survival was significantly different (p = 0.0001, log-rank Mantel-Cox), with increased mortality in GABA_A_R #113-115 IgG high dose (n = 7) and #113-115 Fab high dose (n = 5) groups in comparison to #113-115 IgG low dose (n = 7) and control IgG high dose (n = 5) groups (ANOVA, posthoc Tukey’s multiple comparisons tests; all p ≤ 0.001). (e-h) Quantifications of the indicated (e-f) total and and (g-h) surface proteins from murine brain homogenates after cerebroventricular mAb infusion as analyzed by Western blotting and normalized to mean of control IgG group. Error bars indicate SEM; n = 10 hemispheres per group.

### GABA_A_R mAbs caused spontaneous seizures *in vivo* and spontaneous epileptic activity *ex vivo*

The mAb-induced *in vivo* phenotype suggested hyperexcitability and seizures. We used a wireless EEG system to further evaluate the influence of GABA_A_R mAbs on electric activity at lower concentrations not causing mortality. We examined twelve male Wistar P21 rats that received cerebroventricular infusion of GABA_A_R mAbs or controls and were concurrently implanted with EEG transmitters to record ictal events (Fig. 6a). The EEG coastline length was significantly higher in the GABA_A_R mAb infused animals, indicating increased epileptiform activity (Fig. 6b). For the application of #113-115 and also of α1γ2-dependent #113-175, this correlated with an increase of ictal events as detected by the automated seizure detection program (Fig. 6c, d). The EEG of the GABA_A_R mAb animals showed significantly higher power in all the power band ranges (Fig. 6e).

**Fig. 6.**
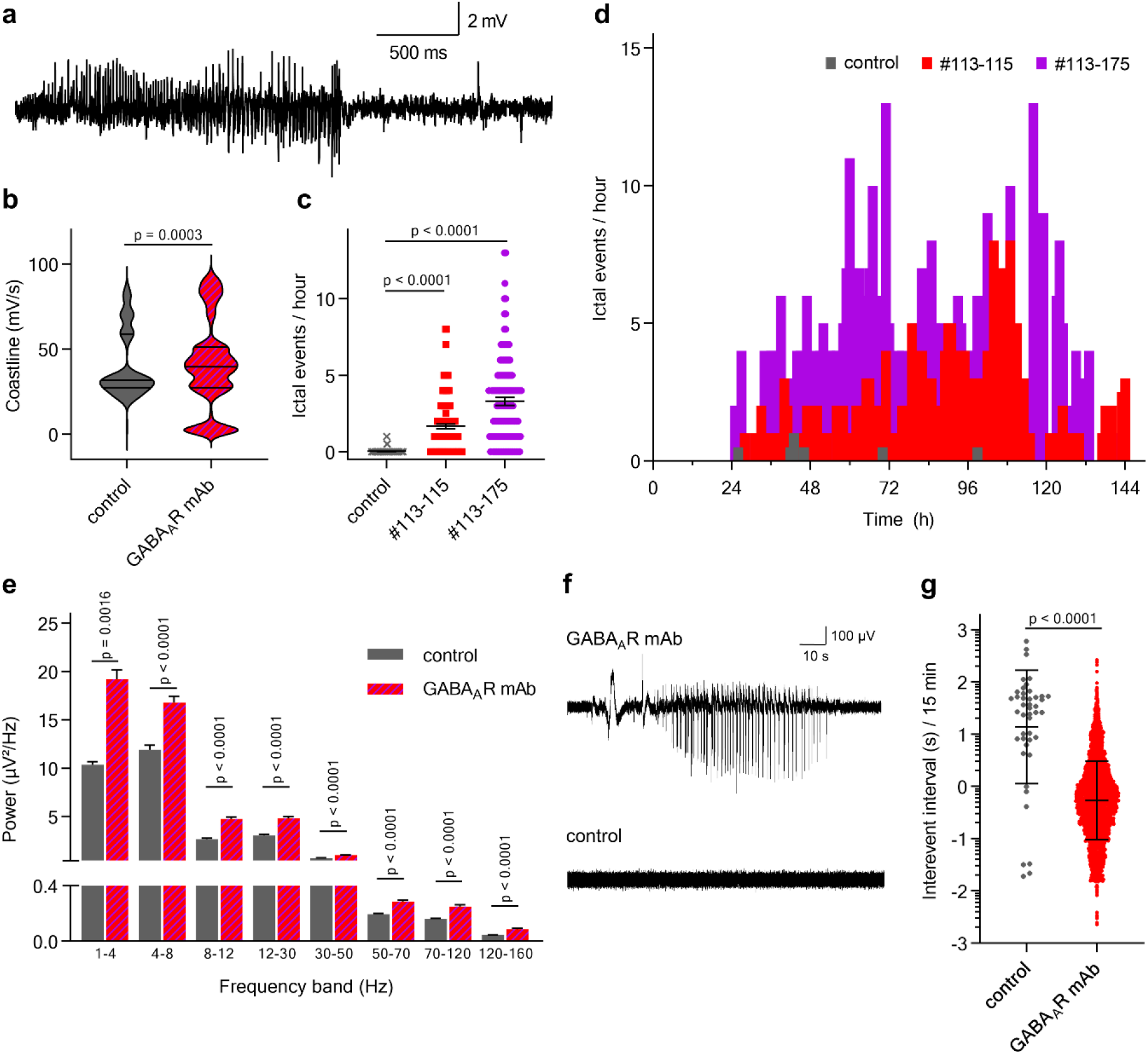
GABA_A_R mAbs caused spontaneous seizures *in vivo* and spontaneous epileptic activity *ex vivo*. (a) Representative EEG of an ictal event recorded from a CA3 depth electrode of a Wistar rat receiving GABA_A_R mAb infusion. (b) Comparison of hourly averages of coastline length between GABA_A_R mAb (#113-115 n = 3, #113-175 n = 3) and control (n = 6) infused animals (Mann-Whitney). Horizonzal lines indicate mean and quartiles. (c) Comparison of the ictal event counts between animals during infusion with GABA_A_R mAb or control as analyzed from Neuroarchiver seizure detection software (Mann-Whitney). Animal numbers as in (b). Bars indicate mean median ± SEM. (d) Distribution of the detected ictal event over time. (e) Comparison of hourly EEG power averages throughout 7-day recordings during GABA_A_R mAb or control infusion (Mann-Whitney). Animal numbers as in (c). (f) Local field potential recording from the CA3 region in a sagittal brain slice from a rat after GABA_A_R mAb infusion (upper trace) showing spontaneous ictal activity *ex vivo*, not seen in controls (lower trace). (g) Comparison of interevent intervals of spontaneous ictal activity from post-mortem acute brain slice recordings from rats after GABA_A_R mAb (#113-115 n = 4, #113-175 n = 3) or control (n = 8) infusion (non-parametric, unpaired Mann-Whitney test). Bars indicate mean ± SEM.

After the completion of infusion with GABA_A_R or control mAbs (day 7-8), acute sagittal brain slices were prepared for local field potential recordings to assess for spontaneous epileptic activity from electrodes placed in areas CA3 and CA1 of the hippocampus. There were significantly higher numbers of spontaneous epileptic events during recordings from the GABA_A_R mAb infused animals compared to controls (Fig. 6f, g). This reaffirms the epileptic activity seen *in vivo* from depth electrodes within the hippocampus of the GABA_A_R mAb infused animals.

## Discussion

Here, we provide insight into the CSF antibody repertoire of acute GABA_A_R encephalitis. Using a recombinant single cell cloning approach, we isolated five monoclonal human GABA_A_R autoantibodies and showed that they recognize GABA_A_Rs *in vitro* and *in vivo*, induce electrophysiological effects independent of receptor internalization and cause a severe encephalitic phenotype in two rodent models. Unlike human serum or CSF containing polyclonal autoantibodies against GABA_A_R and potentially other targets, the mAbs allowed the detailed analysis of antibody sequences, epitope mapping and autoantibody-specific pathogenic functional effects, resulting in a number of novel findings.

The main epitope of all five mAbs was the α1 receptor subunit, as suggested in previous studies^8, 9^. However, one mAb independently co-detected the γ2 subunit, another bound to a shared epitope requiring both α1 and γ2 receptor subunits. Importantly, the latter mAb therefore cannot be detected in current commercial cell-based assays which use α1β3 GABA_A_R, with the potential implication that routine diagnostics in acute encephalitis may underestimate disease-relevant GABA_A_R antibody titers or even overlook affected patients, in case patients harbor only antibodies directed to γ2 receptor subunits in their polyclonal antibody response. The target epitopes of some GABA_A_R mAbs were identical or in close proximity, as shown in our antibody competition assays and indicated from prior experiments with the limitations of CSF polyclonality^8^. Conversely, the presence of mAb #113-115 did not reduce, but enhanced binding of mAb #113-101 to the GABA_A_R. This suggests a direct conformational change of the receptor upon mAb binding, thus potentially adding a new pathogenic principle to the complexity of antibody-induced effects in autoimmune encephalitis. Likewise, conformational effects of disease-relevant mAbs may in some cases stabilize the receptor in a specific activation state, similar to nanobodies which in this way allowed first structure crystallography of G protein-coupled receptors complexed with their G protein^17^.

Our functional investigations *in vitro* and *in vivo* showed the pathogenicity of the two selected GABA_A_R mAbs #113-115 and α1γ2-dependent #113-175. Cerebroventricular infusion of both mAbs induced a severe phenotype with catatonia, seizures and increased mortality, similar to encephalitis patients and in line with epileptic encephalopathies in GABA_A_R mutation carriers^3–6^. However, electrophysiological recordings from autaptic neuronal cultures revealed a reduction of GABAergic currents for mAb #113-115, but not for α1γ2-dependent #113-175. The observed increase of epileptiform activity *in vivo* after #113-175 infusion may possibly be mediated via network effects and thus not detectable in the autaptic *in vitro* model. For both mAbs we did not observe receptor internalization, a previously highlighted mechanism^7, 9^. Possible explanations for the different findings include the use of patient samples containing polyclonal antibodies with undetermined additional specificities in previous studies and the presence of GABA_A_R β3 subunit antibodies not assessed in our study. For #113-115, the electrophysiological findings and missing internalization together with the suggested change of receptor conformation indicate direct functional effects upon mAb binding. Possible mechanisms include stabilization of the receptor in a desensitized state similar to the GABA_A_R agonist benzamidine^18^, allosteric modulation of the physiological GABA binding affinity and/or neurotransmission efficacy, and orthosteric GABA antagonism. Pathogenic mechanisms beyond internalization alone had been suggested by previous results^7^, showing a frequency reduction of IPSCs after GABA_A_R antibody application. Furthermore, our additional *in vivo* experiments using monovalent GABA_A_R Fab fragments not only replicated the severe mAb-induced phenotype, but also supported the concept of direct effects of autoantibody-autoantigen binding, similar to recent studies in which Fab fragments of glycine receptor autoantibodies specifically reduced glycinergic currents^19^. In both cases, Fab fragment experiments prove mAb pathogenicity independent of the integrity of the whole antibody; thereby exclude the dependency of the antibody’s bivalence and Fc-mediated effector functions such as antibody-dependent cell-mediated cytotoxicity, antibody dependent cellular phagocytosis and complement dependent cytotoxicity.

GABA_A_R encephalitis leads to frequent seizures including epilepsia partialis continua and treatment-refractory status epilepticus^8, 10^, which was the predominant phenotype also in our animal models. Using wireless EEG in living animals we could quantify epileptiform activity, including extended measures such as higher coastline length^20^ and increased power in the lower frequency range (1-4 Hz), consistent with EEG changes in human encephalitis patients^21^. Interestingly, in a pharmacologically induced rodent model of status epilepticus, spontaneous seizures were also associated with increased gamma, theta and delta powers in the power spectrum similar to our GABA_A_R mAb model^22^.

Similar to our findings in the related NMDAR encephalitis^12^, we found a broad spectrum of CNS auto-reactivity beyond GABA_A_Rs. These mAbs were reactive to surface and intracellular epitopes on neurons, endothelium and choroid plexus. For example, we identified one anti-Homer-3 antibody, a known target protein in some patients with autoimmune cerebellitis and ataxia^23^. Ongoing attempts to identify other auto-antigens using immunoprecipitation/mass spectrometry, phage display and protein arrays will likely disclose their relevance for additional functional effects, the clinical phenotype and their potential as diagnostic markers. It is tempting to speculate that the diverse non-GABA_A_R mAbs targeting choroid plexus or blood vessels (e.g. mAb #113-201) may contribute to blood-brain barrier dysfunction and thus facilitate entrance of further antibodies and immune cells into the CNS, similar to the role of GRP78 autoantibodies in neuromyelitis optica^24^.

The present study confirmed the importance of isolation and detailed characterization of disease-specific human mAbs from the CSF of patients with encephalitis to foster the comprehensive understanding of humoral CNS autoimmunity. Patient-derived recombinant mAbs represent a useful research tool for multiple purposes and thereby allow the detailed investigation of disease mechanisms and the translation into animal models to recapitulate the clinical phenotype. Most importantly though, retracing the molecular mechanisms of human antibody pathogenicity will provide a refined biological view on some important clinical conditions such as isolated psychosis or catatonia, which can occur with GABA_A_R antibodies^9, 25^. This knowledge will help to reduce stigmatization associated with such psychiatric conditions by understanding them as autoimmune disorders that may require appropriate immunotherapy as a causative treatment in addition to antipsychotics or psychotherapy. Simultaneously, the mAbs are the starting point for the development of novel diagnostics and highly selective immunotherapies for the growing number of patients with antibody-mediated diseases.

## Methods

### Patient sample handling

The index patients’ parents have given written informed consent and analyses were approved by the Charité University Hospital Institutional Review Board.

Four milliliter of CSF were collected during acute phase of encephalitis^14^ and immediately processed for cell pellet cryopreservation, therefore centrifuged for 10 minutes at 400x g, supernatant stored at −80°C and the pellet resuspended in 500 μl of 10% dimethyl sulfoxide, 45% fetal calf serum, 45% RPMI medium before freezing at −80°C. Plasmapheresis eluate was collected six days after lumbar puncture and stored at −80°C.

We used fluorescence-activated cell sorting to isolate single CD138^+^ ASCs, CD20^+^CD27^+^ MBCs and CD20^+^CD27^−^ NMBCs form pre-selected viable CD3^−^CD14^−^CD16^−^DAPI^−^ lymphocytes into 96-well PCR plates. The following antibodies were applied: anti-CD3-FITC (1:25, Miltenyi Biotec, #130-098-162), anti-CD14-FITC (1:25, Miltenyi Biotec, #130-098-063), anti-CD16-FITC (1:25, Miltenyi Biotec, #130-098-099), anti-CD20-PerCP-Vio700 (1:50, Miltenyi Biotec, #130-100-435), anti-CD27-APC-Vio770 (1:12.5, Miltenyi Biotec, #130-098-605) and anti-CD138-PE (1:50, Miltenyi, #130-098-122).

### Generation of recombinant human monoclonal antibodies

From single cell cDNA immunoglobulin genes encoding for variable domains of heavy and light chains were amplified, sequenced and cloned into expression vectors containing the respective constant Ig domains. Sequence analysis data was confirmed by novel custom BASE software^26^. Human embryonic kidney cells (HEK293T) were transiently transfected with an Ig vector pair, mAb containing supernatant harvested and Ig concentrations determined, all following our established protocols^12, 13^.

For reactivity screenings, supernatants were used if concentration was above 2 μg/ml. For detailed characterization of reactivity probabilities and functional assays, cell supernatants were purified using Protein G Sepharose beads, as described before^12^. For Fab fragment synthesis (inVivo BioTech, Henningsdorf, Germany) the IgG1 heavy chain vector was modified by the deletion of the part encoding for the Fc-domains CH2 and CH3 in exchange for a FLAG-Tag and a His-Tag following the amino acids PKSCDKTH of the hinge region. Fab-fragments were purified using immobilized metal affinity chromatography (IMAC).

### Primary neocortical cell cultures

Primary P0-P2 neocortical cultures were prepared from WT Wistar rats, as previously described^27^. Briefly, dissected neocortex tissue was dissociated with Papain for 25 min (1.5 mg/ml; Merck) before trituration in bovine serum albumin (10 mg/ml; Merck). Cells were then resuspended in Neurobasal A medium (supplemented with 1x B27, 1x Glutamax, and 100 U/ml Penicillin-Streptomycin; Thermo Fisher). Dissociated cells were grown in either 24-well or 6-well cell culture plates, coated for 1 hour with Poly-L-lysine hydrobromide (20 μg/ml; Merck). For imaging with the luminescence plate reader, cells were plated in 500 μl droplets @ 400 cells / μl (total: 2×105 cells per well of a 24-well plate). For western blot analysis, cells were plated in 2 ml droplets @ 500 cells / μl (total: 1×106 cells per well of a 6-well plate). Cultures were grown in humidified conditions at 37°C and 5% CO_2_. Cells were cultured until 20-22 days in vitro (DIV) before antibody treatment or fixation.

### Immunohistochemistry

For reactivity screening, recombinant monoclonal antibodies were stained on 20 μm sagittal unfixed mouse brain sections mounted on glass slides. After thawing, tissue was rinsed with PBS, and then blocking solution (PBS supplemented with 2% Bovine Serum Albumin (Roth) and 5% Normal Goat Serum (Abcam)) was applied for 1 hour at room temperature. As primary antibodies, undiluted HEK293T cell supernatants containing recombinant mAbs were incubated overnight at 4°C. After washing three times with PBS, Alexa Fluor 488-conjugated goat anti-human IgG (1:1,000, Dianova, #109-545-003) diluted in blocking solution was added for 2 hours at room temperature, before an additional three time washing and mounting using DAPI-containing Fluoroshield.

For co-stainings, purified recombinant human monoclonal antibodies were used at 5 μg/ml, tissue was fixed with 4% paraformaldehyde (PFA) and stained following the same protocol, but using blocking solution supplemented with 0.1% Triton X-100 (Chemsolute). For co-stainings commercial antibodies rabbit anti-GABA_A_R α1 (1:250, Abcam, #ab33299), guinea pig anti-VGAT (1:500, Synaptic Systems, #131004), chicken anti-MAP2 (1:1,000, Thermo Fisher, #PA1-16751), rabbit anti-MAP2 (1:1,000, Merck, #AB5622), Alexa Fluor 568-conjugated goat anti-chicken IgG (1:500, Invitrogen, #AB_2534098), Alexa Fluor 647-conjugated goat anti-guinea pig IgG (1:1,000, Invitrogen, #A21450), Alexa Fluor 594-conjugated goat anti-rabbit IgG (1:1,000, Dianova, #111-585-003) were used and stainings examined under an inverted fluorescence microscope (Olympus CKX41, Leica DMI6000) or for confocal and large-scale tiling images taken with a cooled EMCCD camera (Rolera-MGi PLUS, QImaging).

### Cell-based assays

HEK293T cells were cultured on Poly-L-lysine coated cover slips, transiently co-transfected with plasmids (kindly provided from Prof. Kneussel from the Center of Molecular Neurobiology in Hamburg, Germany) to express rat α1β3 or α1β3γ2 GABA_A_R and 48 hours later used for live-cell immunostainings following our described protocol, with reduced incubation time for primary antibodies to 1 hour at room temperature, followed by PFA fixation. For neuronal stainings, DIV 20-22 live wildtype neocortical rat neurons were stained with human mAbs for 1 hour at 4°C, washed once before fixation with methanol for 3 minutes at −20°C and then co-stained as described above.

GABA_A_R-negative mAbs with intense tissue reactivity were screened for established neural antigens using commercial panel tests (Euroimmun AG, Lübeck, Germany).

### GABA_A_R subunit reactivity screening

Cell-based binding assay for GABA_A_R was described previously.7 In this study, we additionally cloned cDNAs of rat GABA_A_R α3 (NM_017069), α4 (NM_080587) and β2 (NM_012957) from rat brain total RNA by RT-PCR. COS7 cells were transfected with the indicated GABA_A_R subunits. At 24 hour after transfection, the cells were fixed with 2% PFA at room temperature for 20 minutes and blocked with PBS containing 10 mg/ml BSA for 15 minutes. The fixed cells were incubated with patient derived mAbs (5 μg/ml), followed by staining with the Cy3-conjugated secondary antibody. Then, the cells were permeabilized with 0.1% Triton X-100 for 10 min, blocked with PBS containing 10 mg/ml BSA, and incubated with antibodies to individual GABA_A_R subunits (underlined subunit in the illustration of Fig. 3a and Supplementary Fig. 5), followed by staining with the Alexa488-conjugated secondary antibody. We confirmed that any antibodies did not bind to untransfected cells that did not express the GABA_A_R subunits through distinguishing untransfected cells with Hoechst dye (33342, Thermo Fisher Scientific) nucleic acid staining. images were captured with a system (LSM5 Exciter; Carl Zeiss) equipped with a Plan Apochromat 63x/1.40 NA oil immersion objective lens.

The antibodies used in this screening included: rabbit polyclonal antibodies to GABA_A_R α2 (1:250, Rockland Immunochemicals, Inc. #600-401-D45), α3 (1:250, extracellular epitope; Synaptic Systems #224 303), α5 (1:250, Millipore #AB9678), β3 (1:250, Abcam #ab4046), and γ2 (1:250, extracellular epitope; Synaptic Systems #224 003); guinea pig polyclonal antibodies to GABAA receptor β2 (1:250, Synaptic Systems #224 805); and mouse monoclonal antibodies to GABAA receptor α1 (1:250, NeuroMab #75-136), β1 (1:250, NeuroMab #75-137), and α4 (1:250, NeuroMab 73-383).

The identities of human and rat GABA_A_R α1, α2, α3, α4, α5, β1, β2, β3 and γ2 subunits in their amino acid sequence are 100%, 97%, 95%, 89%, 93%, 98%, 99%, 97% and 99%, respectively; and that of the human and mouse γ2 subunit is 99%, suggesting that the results obtained by using rat or mouse GABA_A_R constructs do not affect the interpretation of our results.

### Quantification of relative binding strength on murine brain sections

Following the protocol above, serial dilutions of purified GABA_A_R mAbs from 50 to 0.002 μg/ml were used for staining of adjacent unfixed murine brain sections. Cerebellar images were recorded with identical settings and analyzed using ImageJ software, version 1.51n (developed by the National Institutes of Health). From each image the MFI was determined from 5 random regions of interest within granular cell layer, secondary antibody MFI subtracted as background, and resulting MFI values scaled to the average MFI values from all five GABA_A_R mAbs at 20 μg/ml^28^. Non-linear regression models (MFI = MFI_max_ * IgG concentration / (Half Max + IgG concentration)) under settings for one site specific binding were generated using GraphPad Prism 8 (GraphPad Software Inc).

### Quantification of relative binding strength via flow cytometry

Following our established protocol^29^, HEK293T cells were transiently co-transfected with plasmids encoding for rat α1, β3 and γ2 subunits of GABA_A_R and enhanced green fluorescent protein (EGFP) or EGFP only and then stained with serial dilutions of purified mAbs. From live single cells with top 30% protein expression (evaluated by EGFP signal) the MFI of the Alexa Fluor 647-conjugated goat anti-human IgG (1:400, Life Technologies) was calculated.

### Competition assay

400 μg of purified GABA_A_R mAbs were concentrated to a volume of 100 μl using Amicon Ultra 100 kDa columns (Merck) and then incubated with 40 μg CruzFluor™ 488 (CF488) succinimidyl ester (Santa Cruz Biotechnology) for 1 hour under rotation in the dark. Unbound fluorophores were washed out using Amicon Ultra 100 kDa columns. IgG concentrations were determined using ELISA and coupling confirmed using UV spectrometry.

CF488-coupled GABA_A_R mAbs were used in serial dilutions to stain unfixed mouse sections and binding was quantified by MFI quantification of CF488 signals to determine individual concentrations for each mAb yielding intensities between detection threshold and signal saturation. At these concentrations (0.4 μg/ml for #113-115, 4 μg/ml, for all others), CF488-coupled mAbs were co-stained with uncoupled mAbs in excess at 50 μg/ml, MFIs quantified and compared to single staining MFIs.

### Quantification of surface protein expression from cultured neurons

To quantify expression of cell surface proteins, we performed state-of-the art biotinylation assays7, 16. In brief, DIV 20-22 rat neocortical neurons were treated with 5 μg/ml of GABA_A_R or control mAb in culture medium for 16 to 18 hours. Neurons were washed, then incubated with EZ-Link Sulfo-NHS-SS-Biotin (Thermo Scientific) for 30 min at 4°C. After quenching unbound biotin, scratched neurons were lysed via sonification in RIPA buffer (150 mM NaCl, 1 mM EDTA, 100 mM Tris HCl, 1% Triton X-100, 1% sodium deoxycholate, 0.1% SDS, pH 7.4, supplemented with protease inhibitor cocktail cOmplete™ Mini (Roche)) for 1 hour at 4°C. Lysates were cleared of debris by centrifugation at 20,000 × g for 20 minutes. The supernatant (whole protein fraction) was incubated with NeutrAvidin UltraLink beads (Thermo Scientific) for 3 hours at 4°C, then eluted (surface proteins fraction). Samples were separated on an 8% gel using SDS-PAGE and analyzed by Western blotting. Images were recorded luminescence-based on ImageQuant^®^ LAS 4000 mini and quantified using ImageQuant TL software (GE Healthcare, version 8.2).

For immunohistochemistry-based receptor quantification we used DIV 20-22 rat neocortical neurons cultured in 24-well plates and applied identical mAb treatments as above. Neurons were stained as live cells with rabbit anti-GABA_A_R α1 antibody (1:1000, Abcam, #ab33299), then washed, fixed with 4% PFA, before application of IRDye^®^ 680RD goat anti-rabbit IgG secondary antibody (1:2000, LI-COR, #925-68071) for 2 hours at room temperature and a final washing with PBS. Plates were recorded in an Odyssey CLx Imaging System (LI-COR) under high-quality resolution settings and analyzed using Image Studio Ver. 5.2 (LI-COR). For each well from a 2000 pixel sized circular region of interest the total pixel intensities were summed, background signals from control wells without primary antibody subtracted and these values scaled to the mean from all wells under untreated condition of this plate.

### Autaptic neurons and electrophysiological recordings

Autaptic murine striatal neurons (DIV 15-18) were incubated with 1 μg/ml human GABA_A_R mAb or control antibody (anti-CD52, Bio-Rad, #HCA175) at 37°C for 24 hours. Data collection from four independent cultures was performed as previously described^30^ with the following alterations. Cells were recorded in standard intra- and extracellular solutions, except for chemically induced NMDA responses, measured in extracellular solution containing 0 mM of Mg2+, 0.2 mM of CaCl2, and 10 μM of glycine. All drugs were bath applied for 1 second except for GABA which was applied for 3 seconds. Currents were normalized per culture to mean of untreated recordings.

### Animals

All animal experiments were performed according to the ARRIVE guidelines and were approved by the Thuringian state authorities (authorization UKJ-17-053) and UK home office guidelines. Male C57BL/6 mice were used at an age of 13-14 weeks and male Wistar rats at post-natal age P21 (weighing 50-58g). Animals were housed in temperature and humidity-controlled conditions on a 12h/12h dark/light cycle and provided with food and water ad libitum.

### Intrathecal osmotic pump infusions

An investigator not involved in the animal experiments randomized mice for assignment to the respective treatment. Animals either received mAb #113-115 as IgG (high dose group: 500 μg over 14 days, 1.5 μg/h, n=7; low dose group: 0.3 μg/h, n=6) or as Fab fragment (equimolar (16.5M) to 1.5 μg/h of IgG, n=5) or control mAb mGO53 as IgG (1.5 μg/h, n=5).

Cerebroventricular infusion with GABA_A_R mAb or control was performed using osmotic pumps (model 1002, Alzet, Cupertino, CA) which were loaded the day before surgery. The following characteristics were used as previously reported^31, 32^: volume 100 μl, flow rate 0.25 μl/hour, and duration 14 days. For surgery, mice were placed in a stereotaxic frame, and a bilateral cannula (model 3280PD-2.0/SP, PlasticsOne) was inserted into the ventricles (coordinates: 0.2 mm posterior and ±1.00 mm lateral from bregma, depth 2.2 mm. Each arm of the cannula was connected to one osmotic pump, which was subcutaneously implanted on the back of the mice (two pumps per animal). After surgery mice were kept single caged and were monitored daily to assess symptoms and survival and were videotaped at representative time points. Mice were sacrificed at day 15. Brain tissue was obtained and frozen in 2-methylbutane.

### Quantification of receptor expression from animals treated via osmotic pumps

For Western blot analysis cryopreserved brain hemisphere from treated mice were thawed and dounced in homogenization buffer (1X PBS, 0.32 M Sucrose, 10 mM HEPES pH 7.4, 2 mM EDTA, 1.6 mM PMSF, supplemented with protease inhibitor cocktail cOmplete™ Mini). Samples were centrifuged at 1,000 x g for 10 minutes. The supernatants (total cell fraction) re-centrifuged at 10,000 x g for 15 minutes and the obtained supernatant finally ultracentrifuged at 100,000 x g for 60 minutes. These pellets were suspended in sample buffer (membrane fraction), separated using SDS-Page and analyzed by Western blotting as above.

### Rat surgery: placement of ventricular catheters, osmotic pumps and wireless EEG transmitters

Osmotic pumps (model 1007D, Alzet) were used for cerebroventricular infusion of GABA_A_R mAbs (volume 100μl, flow rate 0.5 μl/hour, duration 7 days). The day before surgery, two osmotic pumps per animal were prepared by loading with either one mAb or IgG derived from one healthy human aged 35 years. As the initial dose of GABA_A_R #113-115 led to animal death through status epilepticus, applied amounts were reduced in subsequent animals, resulting in ranges from 40-120 μg human mAb (#113-115: n=3, #113-175: n=3, #mGO53: n=4) or 6 mg of polyclonal human IgG (n=2) per animal. The loaded pumps were then connected to polyethylene tubing 69mm-x-1.14mm diameter (C312VT; PlasticsOne) and a double osmotic pump connector intraventricular cannula (328OPD-3.0/SPC; PlasticsOne). Pumps were left overnight in sterile saline solution at 37°C. The next day, under isoflurane anaesthesia, rats were placed in a stereotaxic frame for surgery. The osmotic pumps were placed subcutaneously and attached cannula was inserted into the lateral ventricles (1.5mm lateral, 0.6mm caudal). A subcutaneous pocket was formed over the right flank with a single skin incision and blunt tissue dissection for the transmitter (A3028B-DD subcutaneous transmitters, 90-mm leads, OpenSource Instruments (OSI)), and depth electrode (W-Electrode (SCE-W), OSI) placed in the left hippocampus (CA3, 3.5mm lateral, 3.6mm caudal, depth 2.3mm) with reference electrode implanted in the contralateral skull (3.5mm lateral, 3.6mm caudal). The cannula and skull electrodes were secured with dental cement as previously described^33^. Placement of catheters and electrodes was assessed post mortem during preparation of brain slices for electrophysiology and immunostaining.

### Collection and analysis of EEG data

A custom-built Faraday cage with aerial was used to collect and record EEG data. Transmitter signals were continuously recorded in animals while freely moving using Neuroarchiver software (OSI) and analysed as previously described^33, 34^. In brief, the EEG coastline length was measured as the cumulative absolute difference in voltage between consecutive data points^35^. For automated ictal event detection, video-EEG matching was used to identify ictal EEG events. The Event Classifier (OSI) was then used to classify one second segments of EEG according to several metrics enabling similar events to cluster together when plotted according to metrics. The generated a library of ictal events that allowed fast identification of abnormal EEG events by automated comparison to the library (http://www.opensourceinstruments.com/Electronics/A3018/Seizure_Detection.html). Powerband analysis was carried out using a custom-designed macro.

### Local field potential recordings

At the end of the recording period, 450 μm brain slices were prepared from the GABA_A_R and control mAb infused rats and used for hippocampal local field potential recordings as previously described^36, 37^. Local field potential (LFP) recordings were assessed using Spike2 software (CED) for spontaneous epileptiform activity. Spike2 was used to calculate the root mean square (RMS) amplitude of each recording. Epileptiform activity was classified as an event when it displayed amplitudes greater than four-fold the RMS amplitude, providing the event count, while the time difference between these events provided the interevent interval. Measurements expressed as median (M), interquartile range (Q1-Q3) and min-max values.

### Statistical analysis

All statistical analysis was conducted using GraphPad Prism 8 (GraphPad Software Inc). For sequence analysis numbers of somatic hypermutations were compared using an ordinary one-way ANOVA test followed by posthoc Tukey’s multiple comparisons tests. Amplitudes from autaptic neuron recordings were tested using Kruskal-Wallis, Dunn’s posthoc tests. Protein expression quantifications from Western blotting and immunohistochemistry data were analyzed with ordinary one-way ANOVA tests followed by posthoc Tukey’s multiple comparisons tests. Kaplan-Meier survival analysis was analyzed by log-rank (Mantel-Cox) test followed by ANOVA with posthoc Tukey’s multiple comparisons tests. Coastline length measurements, ictal event counts and power averages from in vivo EEG recordings and interevent intervals of spontaneous ictal activity from post-mortem acute brain slice recordings were analyzed using Mann-Whitney tests.

## Supporting information

All Supplemental Figures and Tables

## Data availability

Raw data were generated at German Center for Neurodegenerative Diseases (DZNE) Berlin, Charité University Medicine Berlin, Aston University Birmingham, University Hospital Jena and National Institute for Physiological Sciences Okazaki. All relevant data supporting the key findings of this study are available within the article and its Supplementary Information files. The mAb sequencing information including raw data files and data derived from custom BASE software analysis of all non-GABA_A_R mAbs from this manuscript have been deposited to Code Ocean (https://codeocean.com/capsule/3514767/). Sequencing information for GABA_A_R mAbs will be added to Code Ocean capsule upon publication of the manuscript and will be shared from the corresponding authors upon reasonable request during review.

## Code availability

The custom software BASE used for Ig sequence analysis is available at https://github.com/automatedSequencing/BASE.

## Acknowledgements

We thank Doreen Brandl, Matthias Sillmann and Stefanie Bandura for excellent technical assistance and Rob Wykes, Jonathan Cornford and Andrea Lieb (University College London, UK) for use of EEG powerband analysis program.

This work was supported by the German Research Foundation (DFG) (FOR3004, GE2519/8-1 and GE2519/9-1 to C.G.; PR 1274/2-1, PR 1274/3-1, and PR 1274/5-1 to H.P.), by the Helmholtz Association (HIL-A03 to H.P.), by the German Federal Ministry of Education and Research (Connect-Generate 01GM1908D to H.P. and 01GM1908D to C.G.), by a BIH-Charité Junior Clinician Scientist Fellowship funded by Charité – Universitätsmedizin Berlin and the Berlin Institute of Health (to J.K. and S.M.R.), by an Epilepsy Research UK Fellowship and Welcome Trust Clinical Research Career Development Fellowship (to S.K.W), by the Schilling foundation (to C.G.), the Interdisciplinary Center of Clinical Research of the Medical Faculty Jena (IZKF; to J.W.), JSPS/MEXT KAKENHI (19H03331, 19K22439 to Y.F.; 20H00459, 19K22548 to M.F.), DAIKO Foundation (to Y.F.), and The Hori Sciences and Arts Foundation (to M.F).

## Author contributions

J.K., S.W., and H.P. designed the study. J.K., S.W., A.v.C., M.L.M., M.N., L.S., S.v.H., E.S.S., J.W., Y.F., M.F., P.T., M.W., S.G. acquired data. J.K., S.W., A.v.C., M.L.M., S.M.R., M.N., H.C.K., J.W., C.G., Y.F., M.F., M.W., S.G., G.W., C.C.G. and H.P. analyzed and interpreted data. J.K., S.W. and H.P. drafted the manuscript. All authors have critically revised and approved the submitted version.

## References

1. Farrant, M. & Nusser, Z. Variations on an inhibitory theme: phasic and tonic activation of GABA(A) receptors. Nat Rev Neurosci 6, 215–229 (2005).

2. Olsen, R.W. & Sieghart, W. International Union of Pharmacology. LXX. Subtypes of gamma-aminobutyric acid(A) receptors: classification on the basis of subunit composition, pharmacology, and function. Update. Pharmacol Rev 60, 243–260 (2008).

3. Hernandez, C.C., et al. Altered inhibitory synapses in de novo GABRA5 and GABRA1 mutations associated with early onset epileptic encephalopathies. Brain 142, 1938–1954 (2019).

4. Wallace, R.H., et al. Mutant GABA(A) receptor gamma2-subunit in childhood absence epilepsy and febrile seizures. Nat Genet 28, 49–52 (2001).

5. Maljevic, S., et al. A mutation in the GABA(A) receptor alpha(1)-subunit is associated with absence epilepsy. Ann Neurol 59, 983–987 (2006).

6. Lachance-Touchette, P., et al. Screening of GABRB3 in French-Canadian families with idiopathic generalized epilepsy. Epilepsia 51, 1894–1897 (2010).

7. Ohkawa, T., et al. Identification and characterization of GABA(A) receptor autoantibodies in autoimmune encephalitis. J Neurosci 34, 8151–8163 (2014).

8. Petit-Pedrol, M., et al. Encephalitis with refractory seizures, status epilepticus, and antibodies to the GABAA receptor: a case series, characterisation of the antigen, and analysis of the effects of antibodies. Lancet Neurol 13, 276–286 (2014).

9. Pettingill, P., et al. Antibodies to GABAA receptor alpha1 and gamma2 subunits: Clinical and serologic characterization. Neurology 84, 1233–1241 (2015).

10. Spatola, M., et al. Investigations in GABAA receptor antibody-associated encephalitis. Neurology 88, 1012–1020 (2017).

11. Bracher, A., et al. An expanded parenchymal CD8+ T cell clone in GABAA receptor encephalitis. Ann Clin Transl Neurol 7, 239–244 (2020).

12. Kreye, J., et al. Human cerebrospinal fluid monoclonal N-methyl-D-aspartate receptor autoantibodies are sufficient for encephalitis pathogenesis. Brain 139, 2641–2652 (2016).

13. Kornau, H.C., et al. Human Cerebrospinal Fluid Monoclonal LGI1 Autoantibodies Increase Neuronal Excitability. Ann Neurol 87, 405–418 (2020).

14. Nikolaus, M., et al. Severe GABAA receptor encephalitis without seizures: A paediatric case successfully treated with early immunomodulation. Eur J Paediatr Neurol 22, 558–562 (2018).

15. Kreye, J., et al. A therapeutic non-self-reactive SARS-CoV-2 antibody protects from lung pathology in a COVID-19 hamster model. Cell 183, 1058–1069 (2020).

16. Hughes, E.G., et al. Cellular and synaptic mechanisms of anti-NMDA receptor encephalitis. J Neurosci 30, 5866–5875 (2010).

17. Rasmussen, S.G., et al. Crystal structure of the beta2 adrenergic receptor-Gs protein complex. Nature 477, 549–555 (2011).

18. Miller, P.S. & Aricescu, A.R. Crystal structure of a human GABAA receptor. Nature 512, 270–275 (2014).

19. Crisp, S.J., et al. Glycine receptor autoantibodies disrupt inhibitory neurotransmission. Brain 142, 3398–3410 (2019).

20. Jones, R.S. & Heinemann, U. Synaptic and intrinsic responses of medical entorhinal cortical cells in normal and magnesium-free medium in vitro. J Neurophysiol 59, 1476–1496 (1988).

21. Symmonds, M., et al. Ion channels in EEG: isolating channel dysfunction in NMDA receptor antibody encephalitis. Brain 141, 1691–1702 (2018).

22. Puttachary, S., et al. Immediate Epileptogenesis after Kainate-Induced Status Epilepticus in C57BL/6J Mice: Evidence from Long Term Continuous Video-EEG Telemetry. PLoS One 10, e0131705 (2015).

23. Höftberger, R., Sabater, L., Ortega, A., Dalmau, J. & Graus, F. Patient with homer-3 antibodies and cerebellitis. JAMA Neurol 70, 506–509 (2013).

24. Shimizu, F., et al. Glucose-regulated protein 78 autoantibody associates with blood-brain barrier disruption in neuromyelitis optica. Sci Transl Med 9 (2017).

25. Pollak, T.A., et al. Autoimmune psychosis: an international consensus on an approach to the diagnosis and management of psychosis of suspected autoimmune origin. Lancet Psychiatry 7, 93–108 (2020).

26. Reincke, S.M., Prüss, H. & Kreye, J. Brain antibody sequence evaluation (BASE): an easy-to-use software for complete data analysis in single cell immunoglobulin cloning. BMC Bioinformatics 21, 446 (2020).

27. Turko, P., Groberman, K., Kaiser, T., Yanagawa, Y. & Vida, I. Primary Cell Culture of Purified GABAergic or Glutamatergic Neurons Established through Fluorescence-activated Cell Sorting. J Vis Exp (2019).

28. Wenke, N.K., et al. N-methyl-D-aspartate receptor dysfunction by unmutated human antibodies against the NR1 subunit. Ann Neurol 85, 771–776 (2019).

29. Ly, L.T., et al. Affinities of human NMDA receptor autoantibodies: implications for disease mechanisms and clinical diagnostics. J Neurol 265, 2625–2632 (2018).

30. Zimmermann, J., Herman, M.A. & Rosenmund, C. Co-release of glutamate and GABA from single vesicles in GABAergic neurons exogenously expressing VGLUT3. Front Synaptic Neurosci 7, 16 (2015).

31. Planaguma, J., et al. Ephrin-B2 prevents N-methyl-D-aspartate receptor antibody effects on memory and neuroplasticity. Ann Neurol 80, 388–400 (2016).

32. Petit-Pedrol, M., et al. LGI1 antibodies alter Kv1.1 and AMPA receptors changing synaptic excitability, plasticity and memory. Brain 141, 3144–3159 (2018).

33. Wright, S., et al. Epileptogenic effects of NMDAR antibodies in a passive transfer mouse model. Brain 138, 3159–3167 (2015).

34. Wykes, R.C., et al. Optogenetic and potassium channel gene therapy in a rodent model of focal neocortical epilepsy. Sci Transl Med 4, 161ra152 (2012).

35. Korn, S.J., Giacchino, J.L., Chamberlin, N.L. & Dingledine, R. Epileptiform burst activity induced by potassium in the hippocampus and its regulation by GABA-mediated inhibition. J Neurophysiol 57, 325–340 (1987).

36. Johnson, N.W., et al. Phase-amplitude coupled persistent theta and gamma oscillations in rat primary motor cortex in vitro. Neuropharmacology 119, 141–156 (2017).

37. Modebadze, T., et al. A Low Mortality, High Morbidity Reduced Intensity Status Epilepticus (RISE) Model of Epilepsy and Epileptogenesis in the Rat. PLoS One 11, e0147265 (2016).

38. Ebert, V., Scholze, P., Fuchs, K. & Sieghart, W. Identification of subunits mediating clustering of GABA(A) receptors by rapsyn. Neurochem Int 34, 453–463 (1999).

