## Supplementary material for "Encephalitis patient derived monoclonal GABA_A_ receptor antibodies cause catatonia and epileptic seizures": All Supplemental Figures and Tables

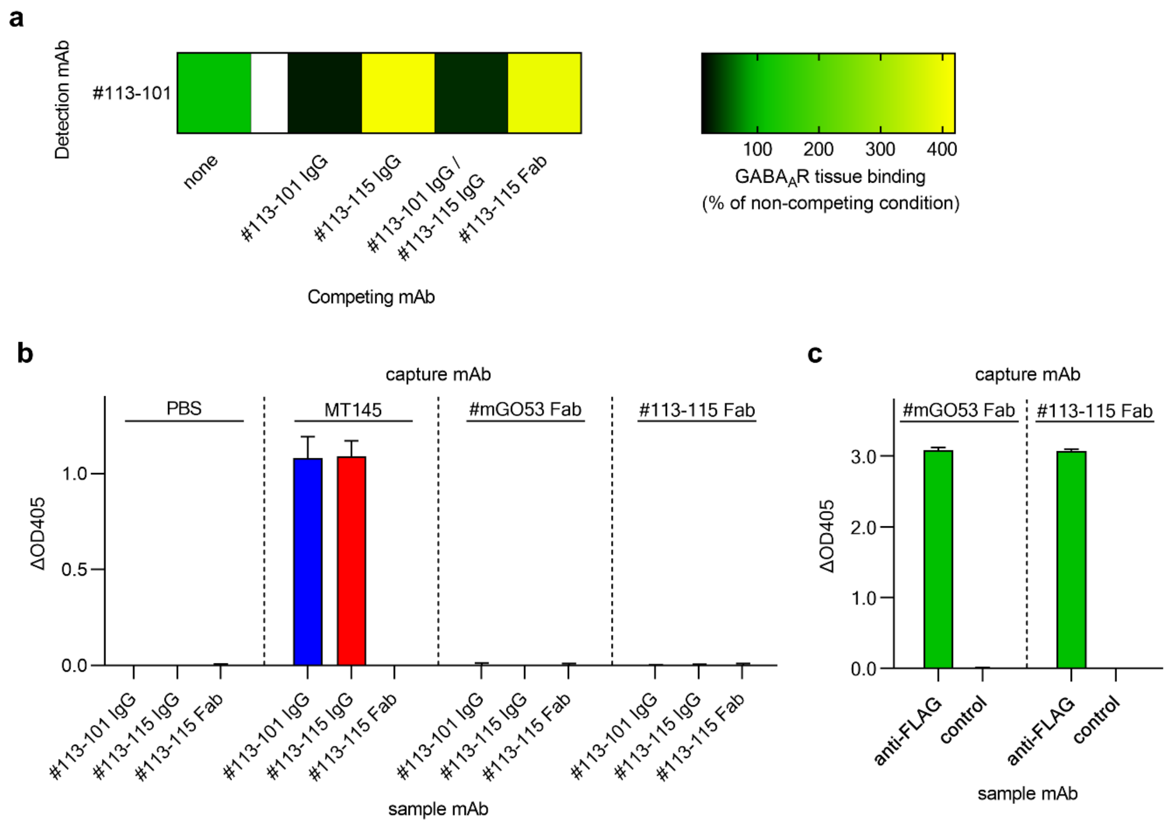


SUPPLEMENTARY INFORMATION

### Encephalitis patient derived monoclonal GABA$_A$ receptor antibodies cause catatonia and epileptic seizures

Jakob Kreye, Sukhvir K. Wright, Adriana van Casteren, Marie-Luise Machule, S. Momsen Reincke, Marc Nikolaus, Laura Stöffler, Scott van Hoof, Elisa Sanchez-Sendin, Hans-Christian Kornau, Max A. Wilson, Stuart Greenhill, Gavin Woodhall, Paul Turko, Imre Vida, Craig C. Garner, Jonathan Wickel, Christian Geis, Yuko Fukata, Masaki Fukata, Harald Prüss

| mAb source cell | | | heavy chain | | | | | light chain | | | | | reactivity | | |
|---|---|---|---|---|---|---|---|---|---|---|---|---|---|---|---|
| **mAb ID** | **pheno-type** | **sub-class** | **IGHV** | **IGHD** | **IGHJ** | **IGH CDR3L** | **IGHV SHM** | **IGK/ IGL** | **IGKV/ IGLV** | **IGKJ/ IGLJ** | **IGK/IGL CDR3L** | **IGKV/IGLV SHM** | **GABA$_A$R CBA** | **brain tissue** | **brain tissue staining pattern** |
| **#113-101** | ASC | IgG1 | 3 | 3 | 4 | 15 | 9 | IGL | 8 | 3 | 10 | 6 | **pos.** | **pos.** | GABA$_A$R pattern |
| **#113-103** | NMBC | IgM | 1 | 3 | 4 | 20 | 15 | IGK | 1 | 1 | 9 | 12 | neg. | neg. | |
| **#113-104** | MBC | IgG1 | 3 | 3 | 4 | 14 | 8 | IGK | 1 | 4 | 9 | 2 | neg. | neg. | |
| **#113-107** | MBC | IgG1 | 3 | 5 | 3 | 15 | 14 | IGK | 3 | 2 | 11 | 11 | neg. | neg. | |
| **#113-109** | NMBC | IgM | 1 | N/A | 4 | 13 | 3 | IGK | 1 | 2 | 9 | 1 | neg. | **pos.** | blood vessels |
| **#113-110** | NMBC | IgM | 3 | 3 | 5 | 19 | 7 | IGL | 1 | 2 | 10 | 2 | neg. | neg. | |
| **#113-111** | NMBC | IgM | 3 | 3 | 3 | 19 | 0 | IGK | 1 | 1 | 9 | 0 | neg. | **pos.** | blood vessels, choroid plexus, basal ganglia |
| **#113-112** | NMBC | IgM | 1 | 3 | 3 | 16 | 3 | IGL | 2 | 1 | 10 | 3 | neg. | neg. | |
| **#113-114** | NMBC | IgM | 3 | 6 | 3 | 15 | 16 | IGL | 2 | 3 | 10 | 10 | neg. | **pos.** | cerebellum (cellular pattern in GCL), hippocampus (neuropil in CA1-3, sparing of MoDG), basal ganglia |
| **#113-115** | ASC | IgG1 | 1 | 3 | 5 | 18 | 12 | IGL | 1 | 2 | 12 | 5 | **pos.** | **pos.** | GABA$_A$R pattern |
| **#113-116** | NMBC | IgG1 | 3 | 4 | 5 | 10 | 10 | IGK | 1 | 2 | 9 | 5 | neg. | neg. | |
| **#113-117** | NMBC | IgM | 3 | 6 | 6 | 21 | 9 | IGL | 1 | 3 | 11 | 6 | neg. | neg. | |
| **#113-118** | NMBC | IgM | 4 | 2 | 6 | 17 | 2 | IGK | 3 | 2 | 8 | 0 | neg. | neg. | |
| **#113-119** | NMBC | IgM | 3 | 4 | 4 | 11 | 3 | IGK | 3 | 2 | 11 | 3 | neg. | neg. | |
| **#113-121** | MBC | IgM | 3 | 7 | 4 | 13 | 11 | IGK | 4 | 2 | 9 | 1 | neg. | neg. | |
| **#113-122** | NMBC | IgM | 1 | N/A | 5 | 12 | 4 | IGL | 1 | 1 | 11 | 4 | neg. | neg. | |
| **#113-123** | NMBC | IgM | 1 | 1 | 4 | 15 | 17 | IGK | 3 | 3 | 9 | 6 | neg. | neg. | |
| **#113-124** | NMBC | IgM | 4 | 1 | 5 | 18 | 1 | IGK | 3 | 4 | 9 | 0 | neg. | neg. | |
| **#113-125** | NMBC | IgM | 3 | 3 | 4 | 13 | 10 | IGK | 3 | 2 | 7 | 0 | neg. | neg. | |
| **#113-126** | MBC | IgG1 | 3 | 3 | 4 | 16 | 15 | IGK | 1 | 1 | 9 | 10 | neg. | **pos.** | fine blood vessels frontal cortex |
| **#113-127** | MBC | IgG1 | 3 | 4 | 4 | 14 | 20 | IGK | 1 | 2 | 9 | 7 | neg. | neg. | |
| **#113-128** | MBC | IgM | 4 | 7 | 5 | 14 | 13 | IGK | 2 | 4 | 9 | 4 | neg. | **pos.** | choroid plexus, blood vessels |
| **#113-130** | NMBC | IgM | 3 | 6 | 6 | 15 | 0 | IGK | 2 | 3 | 9 | 0 | neg. | neg. | |
| **#113-132** | NMBC | IgG1 | 3 | 3 | 4 | 18 | 3 | IGK | 1 | 2 | 9 | 0 | neg. | neg. | |
| **#113-133** | NMBC | IgM | 3 | 1 | 4 | 12 | 13 | IGK | 3 | 4 | 9 | 9 | neg. | neg. | |
| **#113-134** | NMBC | IgM | 3 | N/A | 4 | 12 | 20 | IGK | 3 | 2 | 11 | 11 | neg. | **pos.** | somata in white matter (e.g. corpus callosum) |
| **#113-135** | NMBC | IgG1 | 1 | 1 | 3 | 14 | 15 | IGK | 3 | 1 | 9 | 3 | neg. | **pos.** | cerebellum (PCL, neuropil pattern in GCL and weaker MCL), hippocampus, choroid plexus |
| **#113-136** | NMBC | IgM | 3 | 1 | 4 | 13 | 0 | IGL | 1 | 2 | 13 | 1 | neg. | neg. | |
| **#113-137** | NMBC | IgM | 3 | 1 | 6 | 14 | 8 | IGK | 2 | 1 | 8 | 1 | neg. | neg. | |
| **#113-138** | NMBC | IgM | 4 | 6 | 4 | 11 | 3 | IGK | 1 | 1 | 11 | 6 | neg. | neg. | |
| **#113-139** | NMBC | IgM | 3 | 3 | 2 | 14 | 7 | IGK | 1 | 3 | 9 | 6 | neg. | neg. | |
| **#113-140** | NMBC | IgG1 | 4 | 3 | 4 | 15 | 6 | IGK | 1 | 2 | 9 | 8 | n.a.* | n.a.* | |
| **#113-143** | MBC | IgG1 | 1 | 3 | 6 | 24 | 25 | IGK | 1 | 3 | 9 | 13 | neg. | neg. | |
| **#113-147** | NMBC | IgM | 3 | 1 | 6 | 13 | 0 | IGK | 4 | 1 | 10 | 0 | neg. | neg. | |
| **#113-149** | NMBC | IgM | 3 | 6 | 4 | 17 | 3 | IGK | 1 | 4 | 8 | 2 | neg. | neg. | |
| **#113-159** | NMBC | IgM | 4 | 3 | 6 | 22 | 4 | IGK | 1 | 1 | 9 | 2 | neg. | (pos.) | fine punctuated signal in corpus callosum |
| **#113-160** | NMBC | IgG1 | 4 | 4 | 6 | 18 | 5 | IGK | 3 | 1 | 9 | 7 | neg. | (pos.) | choroid plexus, ependyma cells |
| **#113-162** | NMBC | IgM | 3 | 4 | 4 | 13 | 10 | IGK | 3 | 1 | 9 | 7 | neg. | neg. | |
| **#113-164** | MBC | IgG2 | 3 | 2 | 4 | 14 | 21 | IGK | 1 | 4 | 7 | 32 | neg. | neg. | |
| **#113-166** | NMBC | IgM | 4 | 4 | 4 | 11 | 8 | IGK | 3 | 4 | 9 | 5 | neg. | neg. | |
| **#113-171** | ASC | IgG1 | 4 | 3 | 3 | 23 | 7 | IGK | 1 | 4 | 8 | 8 | neg. | **pos.** | similar to GABA$_A$R pattern |
| **#113-172** | NMBC | IgM | 4 | 5 | 5 | 15 | 0 | IGK | 1 | 2 | 9 | 0 | neg. | neg. | |

| | | | | | | | | | | | | | | | |
|---|---|---|---|---|---|---|---|---|---|---|---|---|---|---|---|
| **#113-174** | NMBC | IgM | 4 | 4 | 6 | 26 | 2 | IGK | 3 | 5 | 8 | 0 | neg. | **pos.** | diffuse ubiquitous staining, pronounced in cerebellum |
| **#113-175** | ASC | IgG1 | 3 | 3 | 4 | 17 | 12 | IGK | 3 | 2 | 10 | 0 | **pos.*** | **pos.** | GABA$_A$R pattern |
| **#113-176** | MBC | IgG3 | 3 | 4 | 3 | 13 | 20 | IGK | 3 | 1 | 8 | 7 | neg. | neg. | |
| **#113-179** | NMBC | IgM | 1 | 3 | 4 | 14 | 0 | IGK | 1 | 3 | 10 | 0 | neg. | (pos) | blood vessels, choroid plexus, ependyma cells |
| **#113-180** | NMBC | IgG3 | 4 | 3 | 5 | 20 | 9 | IGK | 3 | 1 | 9 | 6 | neg. | neg. | |
| **#113-183** | NMBC | IgM | 4 | 1 | 4 | 9 | 4 | IGK | 1 | 1 | 9 | 1 | neg. | neg. | |
| **#113-185** | MBC | IgM | 1 | 2 | 6 | 20 | 3 | IGK | 2 | 2 | 8 | 1 | neg. | neg. | |
| **#113-187** | NMBC | IgG1 | 1 | N/A | 4 | 11 | 8 | IGK | 1 | 2 | 9 | 3 | neg. | neg. | |
| **#113-189** | MBC | IgM | 4 | 1 | 4 | 16 | 17 | IGK | 3 | 1 | 11 | 5 | neg. | neg. | |
| **#113-192** | NMBC | IgM | 4 | 6 | 6 | 17 | 10 | IGK | 3 | 4 | 10 | 6 | neg. | neg. | |
| **#113-198** | ASC | IgG1 | 1 | 3 | 4 | 23 | 7 | IGK | 1 | 1 | 8 | 5 | **pos.** | **pos.** | GABA$_A$R pattern |
| **#113-199** | MBC | IgG1 | 1 | 3 | 4 | 19 | 18 | IGK | 2 | 1 | 9 | 4 | neg. | neg. | |
| **#113-201** | MBC | IgG1 | 4 | 3 | 4 | 15 | 1 | IGK | 1 | 1 | 9 | 11 | **pos.** | **pos.** | GABA$_A$R pattern |
| **#113-202** | NMBC | IgG2 | 3 | 4 | 4 | 6 | 10 | IGK | 2 | 4 | 8 | 5 | neg. | neg. | |
| **#113-204** | NMBC | IgG1 | 4 | 3 | 3 | 21 | 1 | IGK | 4 | 2 | 10 | 0 | neg. | **pos.** | cerebellum (neuropil pattern in MCL), meninges, basalganglia |
| **#113-206** | NMBC | IgM | 3 | 3 | 4 | 16 | 14 | IGK | 1 | 1 | 9 | 14 | neg. | neg. | |
| **#113-207** | MBC | IgM | 3 | 6 | 3 | 17 | 11 | IGK | 1 | 4 | 8 | 5 | neg. | neg. | |
| **#113-208** | NMBC | IgM | 4 | 4 | 4 | 12 | 3 | IGK | 3 | 2 | 9 | 3 | neg. | neg. | |
| **#113-209** | NMBC | IgM | 4 | 4 | 4 | 13 | 0 | IGK | 1 | 2 | 9 | 0 | neg. | neg. | |
| **#113-210** | MBC | IgM | 3 | 3 | 4 | 11 | 3 | IGK | 4 | 4 | 8 | 6 | neg. | **pos.** | cerebellum (cellular pattern in GCL), hippocampus (neuropil in CA1-3), basal ganglia, fibers around corpus callosum |
| **#113-212** | NMBC | IgA1 | 3 | 1 | 3 | 15 | 11 | IGK | 2 | 5 | 9 | 7 | neg. | **pos.** | cerebellum (cellular pattern in PCL), ependyma cells |
| **#113-216** | NMBC | IgG1 | 4 | 4 | 4 | 15 | 3 | IGK | 1 | 1 | 9 | 0 | neg. | **pos.** | choroid plexus, ependyma cells, cortex, cerebellum (pronounced in MCL) |
| **#113-219** | NMBC | IgM | 3 | 3 | 4 | 14 | 15 | IGK | 3 | 4 | 8 | 4 | neg. | neg. | |
| **#113-220** | NMBC | IgG1 | 3 | 3 | 4 | 13 | 0 | IGK | 2 | 1 | 9 | 0 | neg. | **pos.** | cerebellum (cellular pattern in GCL), lateral septal nuclei |
| **#113-221** | MBC | IgG3 | 1 | 6 | 5 | 15 | 27 | IGK | 3 | 1 | 9 | 16 | neg. | neg. | |

**Supplementary Table 1 | Sequence and reactivity data from GABA$_A$R encephalitis cerebrospinal fluid antibody repertoire.** Immunoglobulin sequence features and reactivity characteristics are listed for mAbs with corresponding identifier (mAb ID) isolated from cerebrospinal fluid cells of different phenotypes including antibody secreting cells (ASC), memory B cells (MBC) and non-memory B cells (NMBC). For each mAb the human germline V(D)J gene families of highest sequence identity for the immunoglobulin heavy chain (IGH) and paired light chain of either kappa (IGK) or lambda (IGL) type are noted. The length of complementarity determining region 3 (CDR3L) and the number of somatic hypermutations

(SHM) are given as markers of antibody maturation. Reactivity data is provided from $GABA_AR$ cell-based assays (CBA) and from unbiased reactivity screening on unfixed murine brain tissue sections (pos. = positive, (pos.) = weakly positive, neg. = negative, n.a. = not available as mAb was not expressed, * positive only on CBA with α1β3γ2-$GABA_AR$ but not α1β3-$GABA_AR$). For tissue reactivities staining patterns are described in detail (CA = cornu ammonis, GCL = granule cell layer, MCL = molecular cell layer, PCL = purkinje cell layer, MoDG = molecular layer of the dentate gyrus). Detailed mAb sequencing information including raw data files and data derived from custom BASE software analysis are available on Code Ocean (https://codeocean.com/capsule/3514767/).

| | | #113-101 | #113-115 | #113-175 | #113-198 | #113-201 |
|---|---|---|---|---|---|---|
| **Curve fit** | $R^2$ | 0.96 | 0.89 | 0.91 | 0.97 | 0.90 |
| **$MFI_{max}$** | MFI | 121.4 | 118.3 | 75.8 | 121.4 | 114.7 |
| | Std. Error | 3.3 | 3.6 | 2.2 | 2.6 | 7.7 |
| | 95% CI | 115.0 - 128.0 | 111.3 - 125.3 | 71.5 - 80.3 | 115.9 - 127.1 | 101.4 – 134.5 |
| **Half Max** (= 50% $MFI_{max}$) | Conc. | 2.5 | 0.16 | 0.27 | 0.72 | 6.6 |
| | Std. Error | 0.25 | 0.035 | 0.039 | 0.073 | 1.33 |
| | 95% CI | 2.0 - 3.0 | 0.11 - 0.21 | 0.20 - 0.36 | 0.57 – 0.90 | 4.5 - 10.4 |
| **Relative Ranking** | | 15.7 | 1.0 | 1.7 | 4.6 | 41.7 |

**Supplementary Table 2 | Regression models for $GABA_AR$ mAb binding to unfixed murine brain.** Parameter from best curve fit non-linear regression models of one site specific binding as calculated from mean fluorescence intensity (MFI) values of $GABA_AR$ mAb binding to natively expressed receptors on unfixed murine brain sections in serial dilutions. Regression models are based on the following equation: MFI = $MFI_{max}$ * IgG concentration / (Half Max + IgG concentration). Calculated plateau MFI values are given as $MFI_{max}$, from which concentrations (Conc., in µg/ml) of 50% $MFI_{max}$ (Half Max) are derived. A relative ranking score as indicator of mAb affinity was calculated as ratio of individual Half Max concentration to lowest overall Half Max concentration (0.16 µg/ml from mAb #113-115).

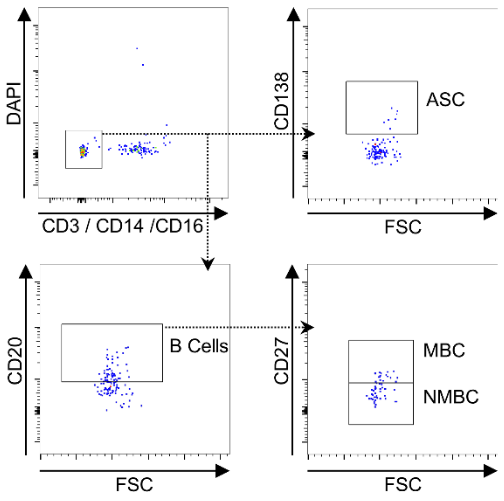


**Supplementary Fig. 1 | Isolation of single cells from cerebrospinal fluid.** Gating strategy in fluorescence-activated cell sorting (FACS) is shown for isolation of CSF single cells for recombinant mAb cloning. $CD3^-CD14^-CD16^-DAPI^-$ lymphocytes (top left) were gated for $CD138^+$ antibody secreting cells (top right) or $CD20^+$ B cells (bottom left), further differentiated into $CD27^+$ memory B cells and $CD27^-$ non-memory B cells (bottom right).

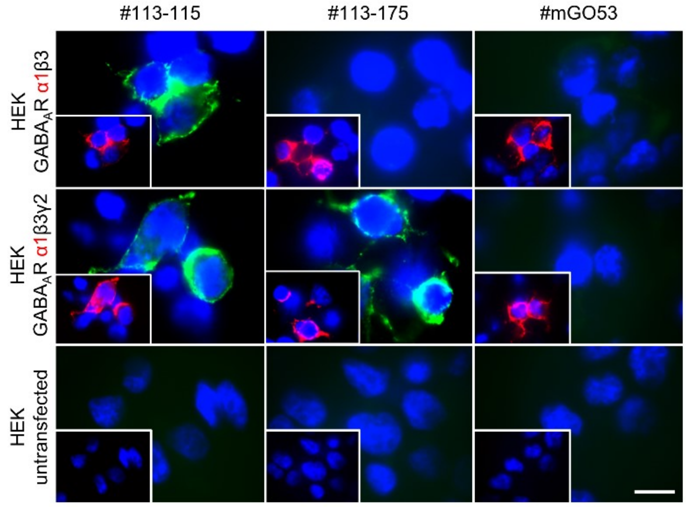


**Supplementary Fig. 2 | Cell-based assays for GABA$_A$R reactivity.** Immunofluorescence stainings of recombinant human mAbs (green, as indicated in column caption) to HEK cells overexpressing the α1β3 or α1β3γ2 subunits of GABA$_A$R or untransfected controls (as indicated in row caption). Co-staining with commercial α1-specific antibody is shown in red and nuclei staining with DAPI in blue. Representative scale bar indicates 20 µm.

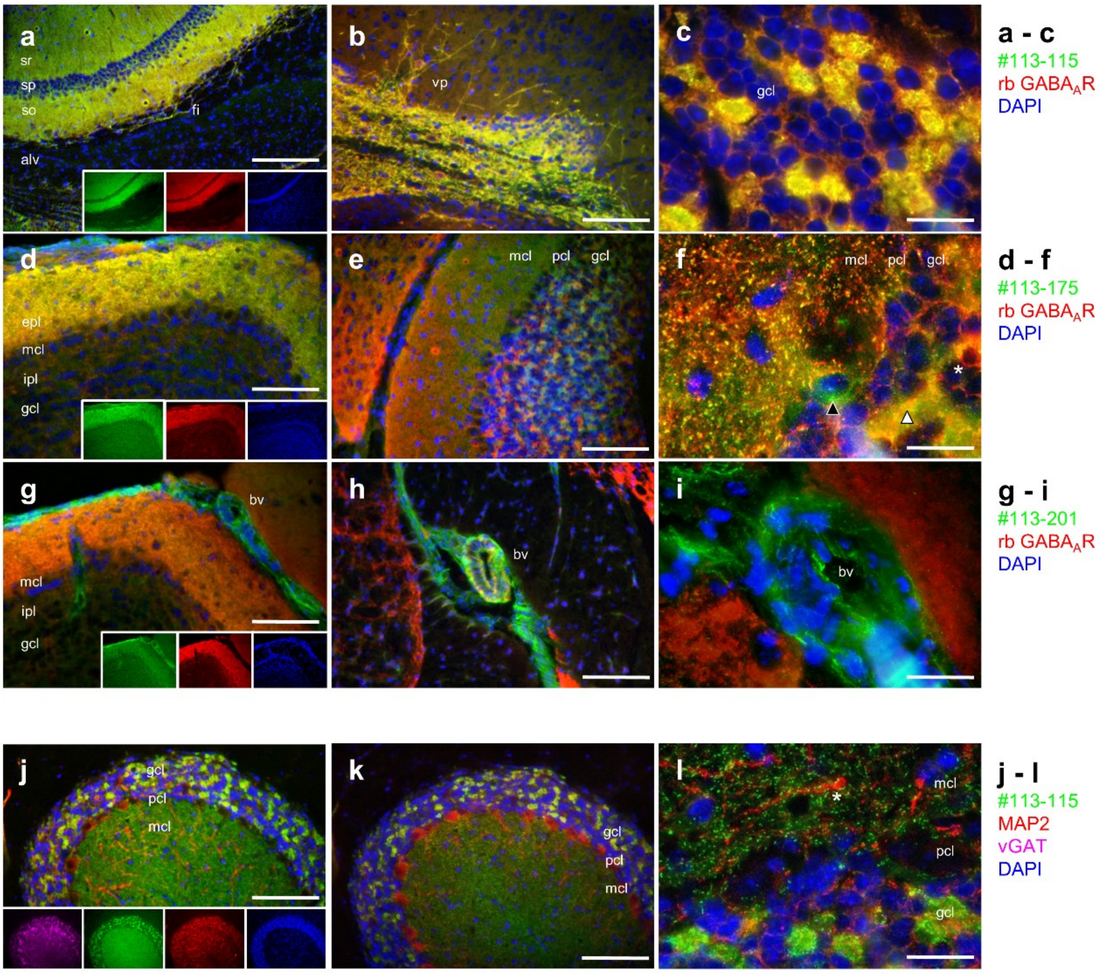


**Supplementary Fig. 3 | Co-localization studies of human GABA$_A$R mAb binding to murine brain tissue with commercial antibodies.** Immunofluorescence stainings of selected human GABA$_A$R mAbs (green) on unfixed murine brain tissue in co-stainings with commercial antibodies (red/pink) and DAPI (blue) as indicated in the row caption on the right. (a) #113-115 intensively stained hippocampal neuropil throughout CA1 and CA2, most pronounced in stratum oriens (so) with hippocampal fimbria (fi) and stratum radiatum (sr), revealing complete overlap with commercial α1-specific antibody binding pattern (red, merge in yellow). (b) Both antibodies showed distinct staining of the ventral pallidum. (c) Higher magnifications in confocal acquisition revealed a somatic staining pattern with complete overlap in the granule cells (gcl) of the cerebellum. (d) α1γ2-dependent #113-175 stained the olfactory bulb, overlapping with commercial α1-specific antibody, most pronounced in the external plexiform layer (epl) and the molecular cell layer (mcl), whereas weaker in the internal plexiform layer (ipl) and the granule cell layer (gcl). (e-f) In the cerebellum, the mcl and gcl pattern uncovered different $GABA_AR^+$ cell populations, some predominantly labelled by #113-175 (green, black arrow), some by commercial antibody (red, white asterisk) and others equally double positive (yellow, white arrow). (g) #113-201 and commercial α1-specific antibody targeted the epl of the olfactory bulb. (h-i) However, #113-201 additionally showed intense binding to blood vessels and choroid plexus, still detectable at dilutions below the $GABA_AR$ pattern detection (not shown). (j) In cerebellar stainings human mAb $GABA_AR$ binding pattern (shown for #113-115) clearly distinguished from microtubule-associated protein 2 (MAP2)-positive dendrites, most pronounced in the mcl, (k) and from vesicular GABA transporter (vGAT)-highlighted somata of the pcl. (l) In magnified confocal images human mAb visualized $GABA_AR$ clusters throughout the mcl and on a subpopulation of cells within the gcl, shown with a reticular MAP2 co-staining. Scale bars indicate 100 µm in left and middle column, 20 µm in right column.

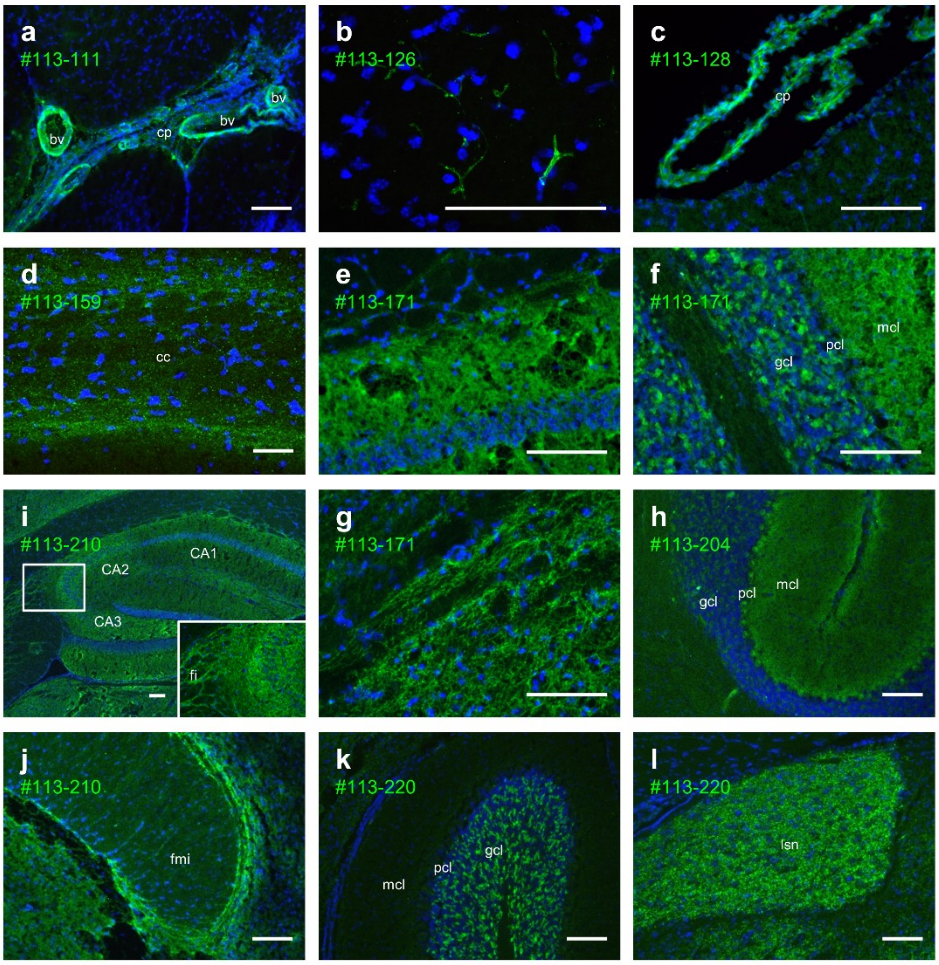


**Supplementary Fig. 4 | Brain tissue reactivity patterns from GABA$_A$R-negative mAbs.** Immunofluorescence stainings of selected human GABA$_A$R-negative mAbs (green, nuclei in blue) from GABA$_A$R encephalitis patient's CSF repertoire with specific binding to unfixed murine brain sections in divergent patterns. Further examples are shown in Fig. 1f-j. (a) Germline mAb #113-111 highlighted large blood vessels (bv) and choroid plexus (cp) tissue. (b) #113-126 stained fine vessels, specifically in frontal cortex. (c) #113-128 targeted choroid plexus. (d) A fine dotted staining pattern in the corpus callosum (cc) was derived from a staining with #113-159. (e-g) #113-171, the only GABA$_A$R-negative mAb derived from an ASC, showed a similar staining pattern as GABA$_A$R mAbs strongly highlighting (e) the hippocampal neuropil as shown in the CA3 region, (f) the granule (GCL) and molecular (MCL) but not Purkinje (PCL) cell layer of the cerebellum and (g) reticular structures in the mesenencephalon. (h) #113-204 was reactive to the MCL and PCL of the cerebellum. (i-j) #113-210 stained intensively hippocampal neuropil, pronounced in the CA2 and CA3 region, fimbria of the hippocampus (fi) and also fine fibers in and around forceps minor (fmi) of corpus callosum. (k) #113-220 targeted selectively granule cells (gcl) in the cerebellum and revealed a punctuated pattern in the lateral septal nuceli (lsn). Scale bars indicate 100 μm.

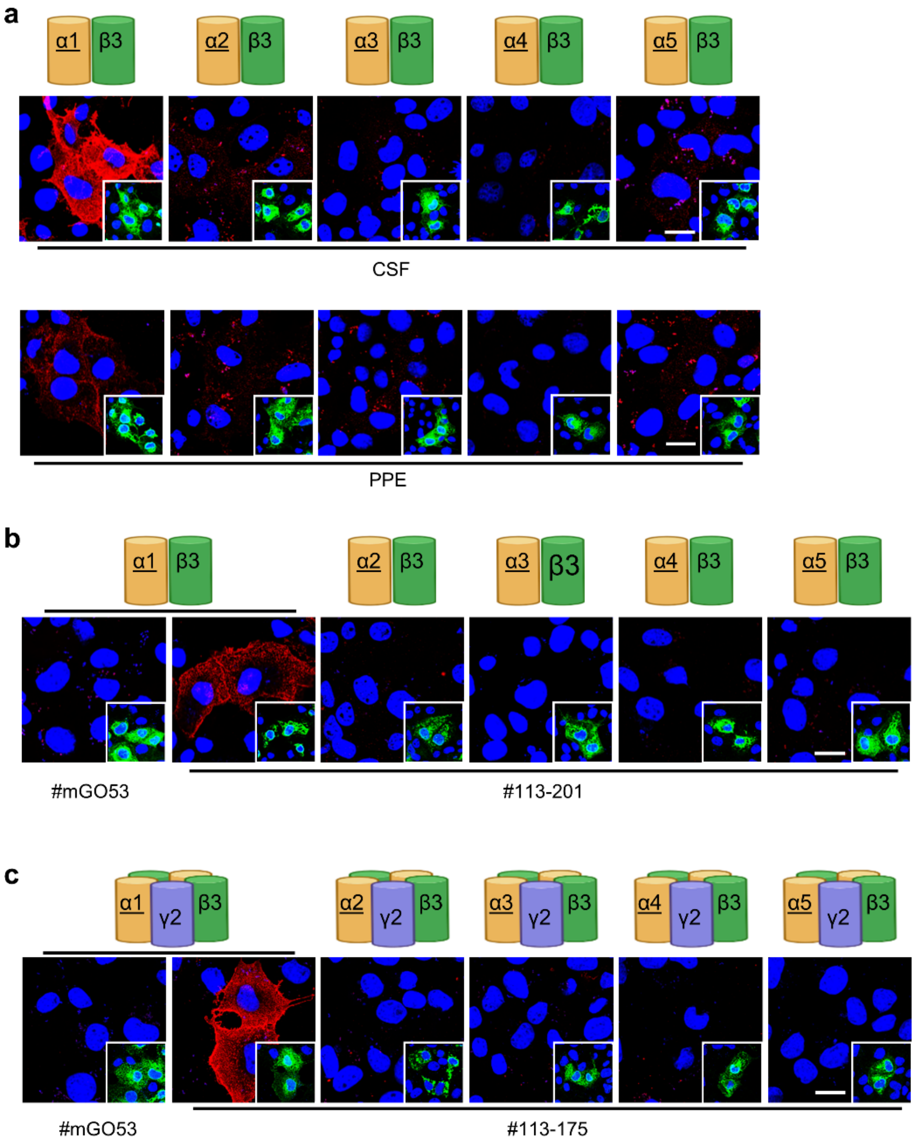


**Supplementary Fig. 5 | Further GABA$_A$R cell-based assays using COS7 cells.** Immunofluorescence stainings as cell-based assays (CBA) using COS7 cells overexpressing individual or multiple GABA$_A$R subunits (as illustrated above) to evaluate subunit specificity of patient's polyclonal samples and derived recombinant mAbs (red, nuclei in blue) in addition to stainings shown in Fig. 3a. Underlined subunits were stained with subunit-specific commercial antibodies (shown in green in image inserts). (a) Patient's polyclonal immunoglobulin samples of cerebrospinal fluid (CSF) and plasmapheresis eluate (PPE) showed CBA binding patterns similar to the mAbs derived from the CSF cells with prominent α1 reactivity (Fig. 3a). Additionally, CSF and PPE antibodies very weakly reacted to COS7 cells expressing α2β3 and α5β3. (b) mAb #113-201 detected the subunits α1 and γ2 independently (Fig. 3a). The α-subunit mediated reactivity is specific to α1, as CBA overexpressing different α-subunits (α2 to α5) in combination with β3 were not detected. Negative control #mGO53 showed no binding. (c) #113-175 specifically detected $GABA_AR$ only when the α1 and γ2 are co-expressed (e.g. α1β3γ2, Fig. 3a), but not in any other combination containing a different α-subunit (α2 to α5) with β3γ2 co-expression. Note that reactivity is absent also when α1β3 or γ2 are expressed alone (Fig. 3a). Representative scale bars indicate 20 μm.

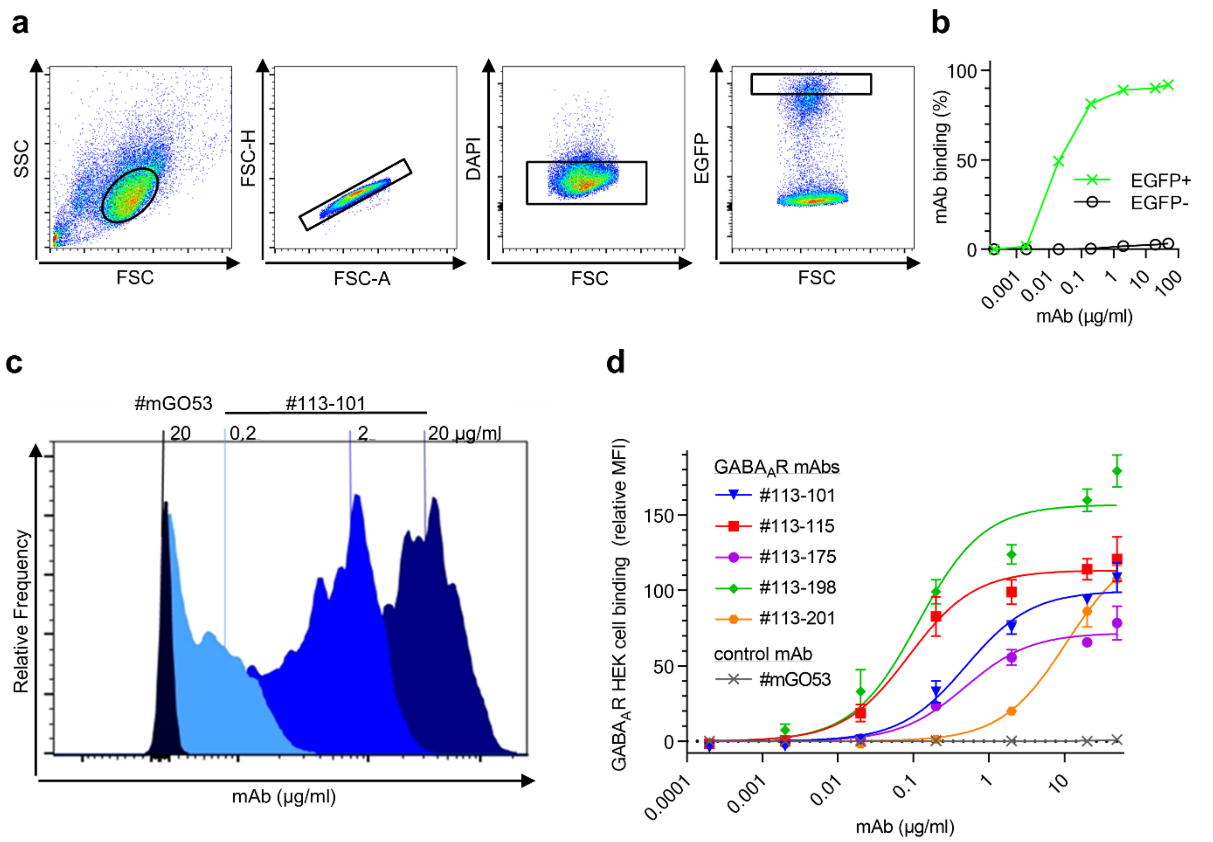


**Supplementary Fig. 6 | Flow cytometry based analysis of GABA$_A$R mAb binding to native receptors.** A flow cytometry approach analyzing HEK cells transfected with α1β3γ2 GABA$_A$R and EGFP was used to complement tissue-based quantification of GABA$_A$R mAb binding to native receptors (Fig. 3b, Supplementary Table 2). (a) Selection of cells for analysis was based on sequential gating on a homogenous cell population in forward (FSC) and sideward scatter (SSC), single cells, live cells (negative for DAPI) and lastly the population of 30% highest enhanced green fluorescent protein (EGFP)-signal as a marker for transfection. (b) Concentration dependent GABA$_A$R mAb #113-115 binding to EGFP-positive and thus GABA$_A$R HEK cells in comparison to EGFP-negative HEK cells. Similar data was obtained from all GABA$_A$R mAbs. (c) Fluorescence histograms of human IgG signal from GABA$_A$R mAb #113-101 (blue) and control mAb #mGO53 (black) stainings at different concentrations are shown with the mean fluorescence intensities (MFI) indicated as vertical lines. (d) MFI values from five independent experiments were used to model binding using non-linear regression models for one site specific binding. Error bars indicate SEM.

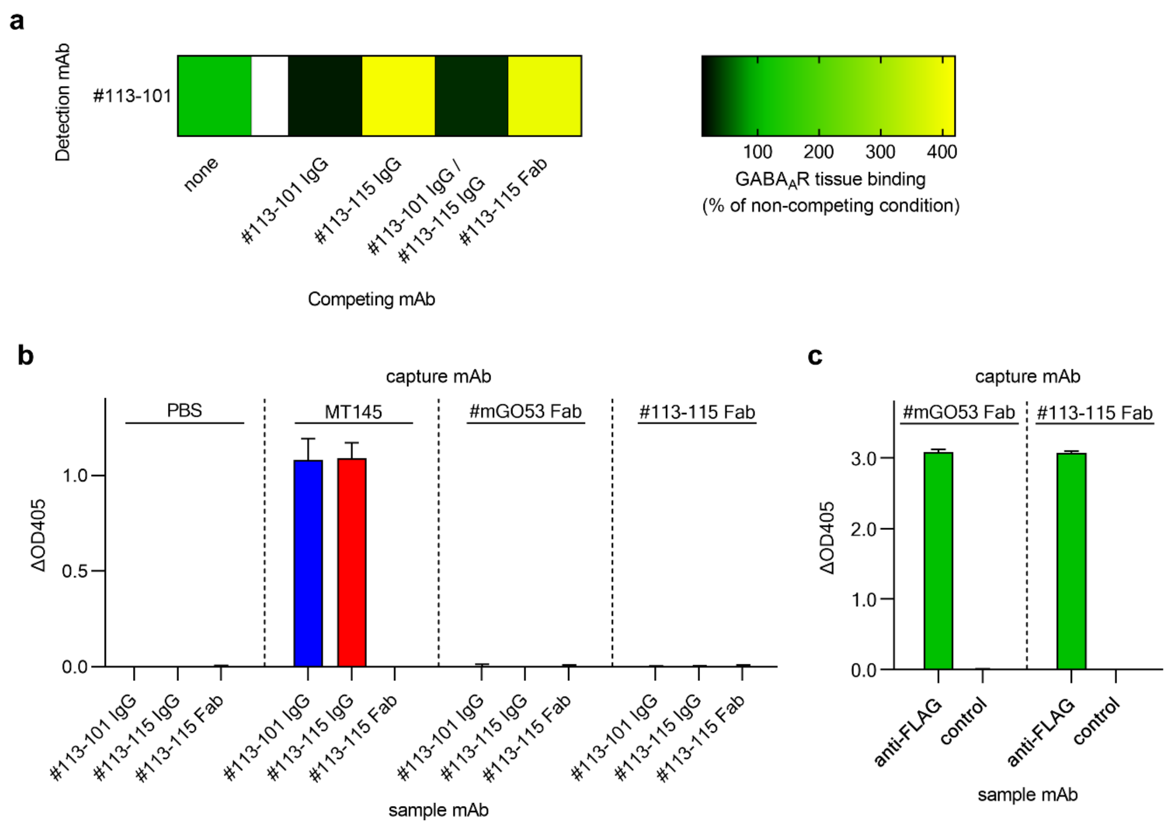


**Supplementary Fig. 7 | Binding interaction of GABA$_A$R mAbs #113-115 and #113-101.** (a) For analysis of competitive binding fluorophore-coupled GABA$_A$R mAb #113-101 (detection mAb) was stained on murine brain tissue in combination with GABA$_A$R mAbs as full IgG or Fab in excess (competing mAb) as indicated. Quantified mean MFIs (see exemplary images in Fig. 3c) as relative values to non-competition conditions are shown as a heat map, each from 30 ROI from 2 independent experiments. Receptor binding competition is visualized in black and signal enhancement in yellow. (b) An ELISA assay was used to exclude binding of #113-101 to the variable domain of the #113-115. High-binding plates were coated with commercial anti-human IgG MT145 or Fab fragments of control mAb #mGO53 or GABA$_A$R mAb #113-115 as indicated in column group titles above. Samples mAbs (human IgG or Fab) were then applied as indicated in column labels below and detected using commercial detection antibody MT78-ALP. Note that MT145 and MT78-ALP are Fc-specific, thus Fabs could not be detected when used as sample mAb. Bars are shown as mean + SD from triplicates of n = 2 experiments. (c) A control ELISA assay was used to confirm successful coating of Fab fragments to high-binding plates. Plates were coated as above before application of commercial mouse anti-FLAG antibody (as sample mAb) for capture by Fab fragments. Bars are shown as mean + SD from triplicates of n = 2 experiments.

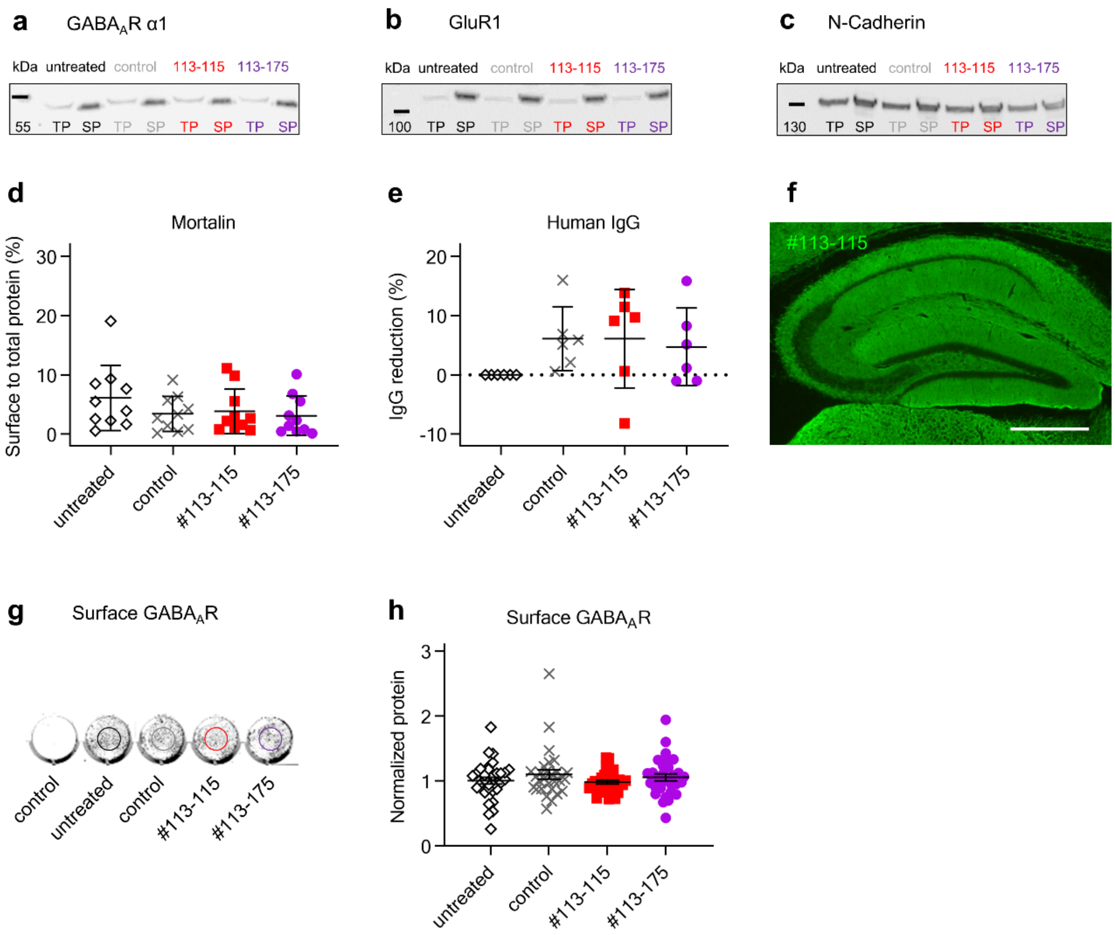


**Supplementary Fig. 8 | Quantification of protein expression from cultured neurons treated with GABA$_A$R mAbs.** (a-c) Representative western blots of total protein (TP) and biotinylated surface protein (SP) samples from neocortical rat neurons after pre-incubation with indicated GABA$_A$R or control mAb #mGO53. Blots were stained with commercial antibodies against the (a) α1-subunit of GABA$_A$R, (b) the GluR1-subunit of AMPA receptors and (c) N-Cadherin. Quantifications of protein levels are shown in Fig. 4g-j. (d) Western blot based quantifications of mortalin levels from the same neocortical rat neuron cultures. Surface to total protein fraction ratios of mortalin, a stricly intracellular protein, indicated no influence of mAb pre-incubation on membrane integrity. Error bars indicate SD; n = 4 separate cultures. (e) Quantification of human IgG from neocortical rat neuron culture media used for Western blotting experiments (a-d and Fig. 4g-j). The reduction of IgG was measured as the relative difference between post-treatment medium and matched source medium samples, which have not been applied to the culture (relative reduction = (concentration $_{source\ medium}$ – concentration $_{post\text{-}treatment\ medium}$) / concentration $_{source\ medium}$). Error bars indicate SD; n = 2 separate cultures. (f) Representative immunofluorescence staining of post-treatment neuronal culture medium containing mAb #113-115 on an unfixed murine brain section confirmed $GABA_AR$ reactivity of the antibody by typical staining pattern, as shown here in hippocampus. Scale bars indicates 500 µm. (g) Representative reader-based immunocytochemistry recordings from neocortical rat neurons after pre-incubation with indicated $GABA_AR$ or control mAbs. Live neurons were stained with a commercial antibody against the α1-subunit of $GABA_AR$ or without primary antibody (control). (h) Quantifications of $GABA_AR$ surface levels from reader-based immunohistochemistry recordings revealing no difference. Error bars indicate SEM; n = 4 separate cultures.

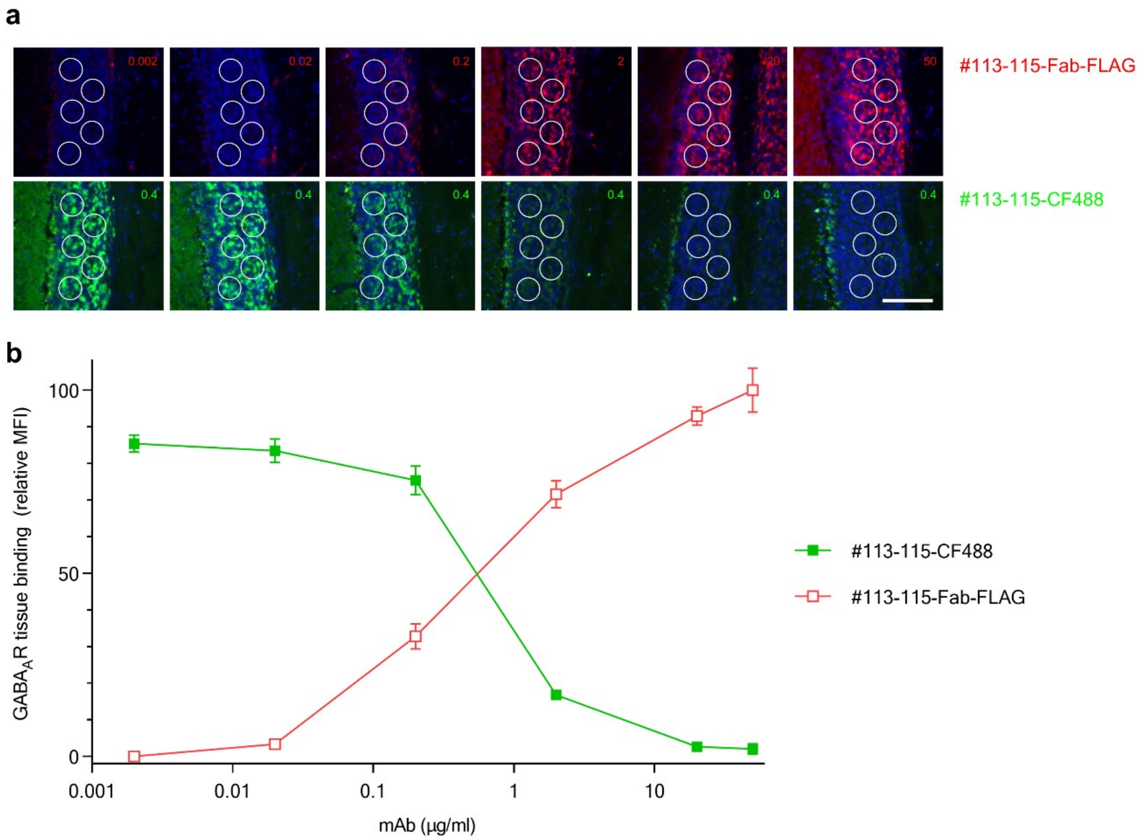


**Supplementary Fig. 9 | Competition of mAb #113-115 Fab and #113-115 IgG for binding to murine brain tissue.** (a) To confirm competitive binding to the identical target GABA$_A$R mAb #113-115 as Fab fragment was stained at serial dilutions as indicated in the presence of fluorophore-coupled #113-115 IgG (#113-115-CF488) at threshold concentration of 0.4 µg/ml (green). Binding of Fab was visualized using an anti-FLAG secondary antibody (red, nuclei in blue). Mean fluorescence intensity (MFI) was measured separately for Fab and IgG binding in identical ROI within granule cells of the cerebellum. Representative scale bar indicating 100 µm. (b) MFI values from 18 ROI of n = 2 independent experiments are shown as mean ± SEM.

#### Supplementary Videos

**Supplementary Video 1 | Catatonia and epileptic seizures in mice after intrathecal infusion of GABA$_A$R mAbs.** (A) Control mouse receiving high-dose control IgG showed normal explorative behaviour 3 days after intrathecal infusion. (B) In contrast, infusion of high-dose GABA$_A$R mAb #113-115 led to severe catatonia and focal epileptic seizures after 3 days. (C) Direct comparison of spontaneous behaviour in mice receiving either GABA$_A$R (left) or control mAb (right). (D) In mice receiving lower doses of GABA$_A$R mAb, catatonia and focal seizures occurred later (day 6) and were less pronounced. (E) Worsening of motor abnormalities at day 10 in animals receiving high-dose GABA$_A$R mAb with hyperexcitability and increased seizure frequency. (F) In contrast, control mice remained healthy and showed normal explorative behaviour after 14 days of intrathecal mAb infusion.
